# Transcriptome analysis of alternative splicing-coupled nonsense-mediated mRNA decay in human cells reveals broad regulatory potential

**DOI:** 10.1101/2020.07.01.183327

**Authors:** Courtney E. French, Gang Wei, James P. B. Lloyd, Zhiqiang Hu, Angela N. Brooks, Steven E. Brenner

## Abstract

To explore the regulatory potential of nonsense-mediated mRNA decay (NMD) coupled with alternative splicing, we globally surveyed the transcripts targeted by this pathway via RNA-Seq analysis of HeLa cells in which NMD had been inhibited. We identified putative NMD-targeted transcripts as those with a termination codon more than 50 nucleotides upstream of an exon-exon junction (premature termination as defined by the ‘50nt rule’) and that significantly increased in abundance upon NMD inhibition. We additionally controlled for potential transcriptional up-regulation by requiring the putative NMD targets to increase in abundance substantially more than the isoforms from the same gene that do not contain a premature termination codon. This resulted in a conservative set of 2,793 transcripts derived from 2,116 genes as physiological NMD targets (9.2% of expressed transcripts and >20% of alternatively spliced genes). Our analysis identified previously inferred unproductive isoforms and numerous heretofore-uncharacterized ones. NMD-targeted transcripts were derived from genes involved in many functional categories, and are particularly enriched for RNA splicing genes as well as for those harboring ultraconserved elements. By investigating the features of all transcripts impacted by NMD, we find that the 50nt rule is a strong predictor of NMD degradation while 3’ UTR length on its own generally has only a small effect in this human cell line. Additionally, thousands more transcripts without a premature termination codon in the main coding sequence contain a uORF and display significantly increased abundance upon NMD inhibition indicating potentially widespread regulation through decay coupled with uORF translation. Our results support that alternative splicing coupled with NMD is a prevalent post-transcriptional mechanism in human cells with broad potential for biological regulation.

## Introduction

Nonsense-mediated mRNA decay (NMD) is a eukaryotic mRNA surveillance pathway with a role in gene regulation. In its surveillance capacity, NMD recognizes premature termination codons (PTCs) generated by nonsense mutations or errors in transcription and splicing and eliminates the aberrant transcripts in order to protect the cell from the production of potentially harmful truncated proteins (Frischmeyer-Guerrerio et al. 2011; He et al. 1993; Isken and Maquat 2007; Kervestin and Jacobson 2012; Lykke-Andersen and Bennett 2014; Pulak and Anderson 1993). Many other transcripts targeted to NMD are generated by highly regulated and evolutionarily conserved splicing events (Lareau, Inada, et al. 2007; Lareau and Brenner 2015; Mudge et al. 2011; Ni et al. 2007; Yan et al. 2015). This coupling of alternative splicing and NMD (AS-NMD) can be used to post-transcriptionally regulate the level of mRNA (also termed Regulated Unproductive Splicing and Translation, or RUST) (He and Jacobson 2015; Kurosaki, Popp, and Maquat 2019; Lewis, Green, and Brenner 2003; Lykke-Andersen and Jensen 2015; Rehwinkel et al. 2005; Schweingruber et al. 2013).

A number of splicing factors are known to auto-regulate their own expression or regulate the expression of other splicing factors through alternative splicing coupled with NMD (Änkö et al. 2012; Dredge and Jensen 2011; Jangi et al. 2014; Jumaa and Nielsen 1997; McGlincy et al. 2010; Pervouchine et al. 2019; Rossbach et al. 2009; Saltzman et al. 2008; Saltzman, Pan, and Blencowe 2011; Spellman, Llorian, and Smith 2007; Sun et al. 2010; Sureau et al. 2001). Many of these NMD-coupled splicing events (including all those in the SR gene family) are associated with a highly conserved or ultraconserved element (Bejerano et al. 2004) of the human genome, indicating a potential role for these elements in the regulation of alternative splicing coupled with NMD (Lareau, Brooks, et al. 2007; Ni et al. 2007). Genes with other functions have also been reported to be regulated via alternative splicing coupled to NMD such as the spermidine/spermine N1-acetyltransferase (SSAT) gene (Hyvönen et al. 2006, 2012) and ribosomal proteins (Cuccurese et al. 2005; Plocik and Guthrie 2012; Russo et al. 2011; Zheng et al. 2012). In addition, NMD, not always coupled to alternative splicing, has been shown to be important for multiple differentiation programs including neuronal, granulocyte, and erythropoiesis differentiation (Bruno et al. 2011; Karam and Wilkinson 2012; Lasalde et al. 2014; Pimentel et al. 2014; Weischenfeldt et al. 2008; Wong et al. 2013), essential for viability via regulation of *GADD45*, a stress and DNA-damage response factor (Nelson et al. 2016), and playing a role in cancer (Fernandes et al. 2019).

Numerous studies have been performed to investigate the components modulating the NMD process, resulting in identification of many NMD factors (Bhattacharya et al. 2000; Deniaud et al. 2015; Gehring et al. 2005; Huntzinger et al. 2008; Ivanov et al. 2008; Kashima et al. 2006; Lloyd et al. 2018; Lykke-Andersen and Bennett 2014; Lykke-Andersen, Shu, and Steitz 2000; Melero et al. 2014; Yamashita et al. 2009). Among those proteins known to mediate the NMD pathway, UPF1 is a crucial and conserved component (Chakrabarti et al. 2011, 2014; Hwang et al. 2010; Lasalde et al. 2014; Nicholson et al. 2014; Okada-Katsuhata et al. 2012). However, other than UPF1, other NMD factors may be exclusive to different branches of the pathway (Metze et al. 2013) and the specific mechanisms of NMD and the defining features of NMD targets are still not completely understood and likely vary across different species (Kerényi et al. 2008; Lloyd 2018). In mammals, a prominent model posits that the presence of an exon-exon junction more than 50 nucleotides downstream of a stop codon induces NMD; this has been termed the “50 nucleotide rule” (Nagy and Maquat 1998; Zhang and Maquat 1997). The exon junction complex is a protein complex deposited near an exon-exon junction during the splicing process and acts as a cellular memory of an intron’s location following splicing. When a ribosome terminates upstream of an exon-exon junction, NMD factors in the exon junction complex recruit UPF1, triggering decay (Chamieh et al. 2008). Long 3’ UTRs have also been found to target transcripts to NMD (Amrani et al. 2004; Balagopal and Beemon 2017; Bao et al. 2016; Bühler et al. 2006; Chakrabarti et al. 2014; Fanourgakis et al. 2016; Ge et al. 2016; Hurt, Robertson, and Burge 2013; Kervestin et al. 2012; Kervestin and Jacobson 2012; Kishor, Fritz, and Hogg 2019; Kishor, Ge, and Hogg 2019; Kurosaki and Maquat 2013; Lareau and Brenner 2015; Nyikó et al. 2017; Singh, Rebbapragada, and Lykke-Andersen 2008; Toma et al. 2015; Wright et al. 2015; Yepiskoposyan et al. 2011). A long 3’ UTR, or “faux” 3’ UTR, has been proposed to trigger NMD in a couple of ways including structural characteristics, such as the long distance between PABP on the poly-A tail and the terminating ribosome, lead to aberrant termination (Behm-Ansmant et al. 2007; Eberle et al. 2008; Kervestin et al. 2012) or that UPF1 binds to 3’ UTRs in a length-dependent manner and this primes the transcripts for NMD (Hogg and Goff 2010; Zünd et al. 2013).

The potential prevalence of NMD-targeted alternative isoforms was first established by searching EST databases for transcripts with a PTC (Lewis et al. 2003). These datasets, however, were generated from cells where NMD was active and the targets should be depleted. There have also been studies that successfully identified NMD-targeted transcripts by inhibiting NMD (often by perturbing UPF1) and using microarray methods (Hansen et al. 2009; Mendell et al. 2004; Pan et al. 2006; Saltzman et al. 2008; Sayani et al. 2008; Wengrod et al. 2013; Wittmann, Hol, and Jäck 2006; Yepiskoposyan et al. 2011). However, microarray analyses generally rely on probes derived from known transcript sequences, again, derived from normal cells where NMD is active. Thus, their capacity to detect native transcripts targeted by NMD is limited, and many novel isoforms were likely missed. High-throughput deep sequencing of the transcriptome and current analytical algorithms now allow for the improved identification and quantification of novel isoforms. These methods have been combined with NMD inhibition and used to investigate the effect of NMD on the transcriptome in mouse (Hurt et al. 2013; Weischenfeldt et al. 2012) and in human (Colombo et al. 2017; Lou et al. 2016; Lykke-Andersen et al. 2014; Tani et al. 2012) and deep sequencing of mouse and human brains has revealed extensive alternative splicing coupled to NMD, even in the presence of active NMD (Yan et al. 2015).

Many questions regarding the prevalence and mechanism of NMD remain open. In this study we used NMD inhibition via knockdown of UPF1 and RNA-Seq analysis in a human cell line (HeLa) in order to discover genes that produce putative unproductive isoforms and investigate the features of transcripts affected by NMD. For both known and novel transcripts, we predicted isoform-specific putative PTCs (by the 50nt rule) and measured isoform-level expression changes, in contrast to other studies that used gene expression and annotated coding sequences. Using isoform-level analyses allows us to then filter out genes that are transcriptionally up-regulated due to secondary effects of the experiment. Even with this more conservative analysis, we found that a substantial fraction of alternatively spliced transcripts are subject to NMD, even in just this single cell type and condition. Furthermore, we were able to investigate the features of NMD-targeted isoforms and found that the presence of an exon-exon junction ≥50nt downstream of a stop codon was a strong predictor of NMD susceptibility, as expected. However, the length of the 3’ UTR on its own had a much smaller effect globally. We also found that NMD targets are enriched for the presence of ultraconserved elements and for splicing regulators and RNA-binding proteins. Additionally, many transcripts with uORFs are sensitive to NMD. Our findings include numerous previously unreported substrates for this process in human cells, implying broad regulatory potential.

## Results

### Over two thousand genes produce transcripts identified as high confidence NMD targets

Transcripts with a premature termination codon (PTC), defined as an early stop codon that triggers NMD, should be stabilized if the NMD pathway is inhibited. To systematically survey NMD-targeted transcripts, we performed strand-specific paired-end RNA-Seq on HeLa cells with inhibited NMD. To inhibit NMD, we used short hairpin RNAs (shRNAs) against the mRNA of NMD factor UPF1 (Bühler et al. 2006; Paillusson et al. 2005). For each of the two biological replicates, a paired control experiment using an shRNA of random sequence was also performed (Bühler et al. 2006). Western blot analysis confirmed that the protein level of UPF1 in the knockdown cells was 6% of that in the control cells and real-time PCR showed that the abundances of the known NMD-targeted isoforms of *SRSF2* and *SRSF6* were substantially increased (4-30x), suggesting that NMD was substantially inhibited (**Figure S1**). The RNA-Seq reads were then aligned to the human reference genome with Tophat (Trapnell, Pachter, and Salzberg 2009) and assembled into known and novel isoforms with Cufflinks (Roberts et al. 2011; Trapnell et al. 2010). After filtering out transcripts with low confidence junctions (see Supplementary Methods), the transcript level abundance was quantified for a set of 131,003 isoforms, 60,976 of which were substantially expressed (**Table S1**).

We then defined a set of high-confidence direct NMD targets by focusing on those transcripts that showed compelling evidence of being NMD targets, by the 50nt rule model and by our experimental data, as well as by controlling for experimental secondary effects. To do this, we first looked for transcripts with a termination codon more than 50 nucleotides upstream of the transcript’s last exon-exon junction. Such a stop codon, which we term a ‘PTC_50nt_’, would be a putative PTC according to the 50nt rule and is expected to trigger NMD. In order to determine which isoforms have a PTC_50nt_, we predicted the coding sequence (CDS) of each isoform for each gene (see Supplementary Methods), resulting in 30,317 expressed CDS-containing isoforms derived from 11,055 genes (**Table S1**). We found that a fifth (6,429) of these transcripts contained a PTC_50nt_.

Direct targets of NMD should also exhibit significantly higher abundance when NMD was inhibited. We found 10,717 CDS-containing transcripts that increased at least 1.5x and had significant differential expression when NMD was inhibited (**Table 1**). These were significantly enriched for PTC_50nt_-containing transcripts (Fisher’s exact test, p < 2.2 × 10^−16^, which is the limit in R). Altogether, 3,832 out of 6,429 PTC_50nt_-containing transcripts (60%) were significantly and 1.5x more abundant in NMD inhibited cells, compared to only 871 (14%) that were similarly less abundant (**Table 1, Figure 1A**). Many other PTC_50nt_-containing transcripts demonstrated increased abundance, but were not statistically significant. Confounding transcriptional or splicing changes and uncertainty in the analysis could explain most of the PTC_50nt_-containing transcripts that do not increase. Compared to transcripts with a normally positioned non-PTC_50nt_ termination codon (an NTC), PTC_50nt_-containing transcripts demonstrated a strong bias towards accumulation when NMD was inhibited (Kolmogorov-Smirov (KS) test, p < 2.2 × 10^−16^). Transcripts with an NTC had similar percentages of significantly increased (29%) and decreased (28%) transcripts (**Figure 1B**), suggesting that they may be affected by secondary effects from the knockdown of the UPF1 protein and are unlikely to have been directly affected by NMD.

**Table 1.** Classification of expressed transcripts with and without a PTC50nt.

|  | Number of isoforms | Increased abundance<sup>†</sup> | Decreased abundance<sup>†</sup> |
| --- | --- | --- | --- |
| All expressed transcripts with a defined coding sequence | 30,317 | 10,717 (35%) | 7,600 (25%) |
| Transcripts with a NTC | 23,888 | 6,885 (29%) | 6,729 (28%) |
| Transcripts with a PTC<sub>50nt</sub> | 6,429 | 3,832 (60%) | 871 (14%) |

<sup>†</sup>Increased and decreased abundance refers to the expression changes in NMD-inhibited cells (compared to control cells) that are significant according to Cuffdiff and >1.5x. The percentage is of the number of transcripts in that category.

**Figure 1.**
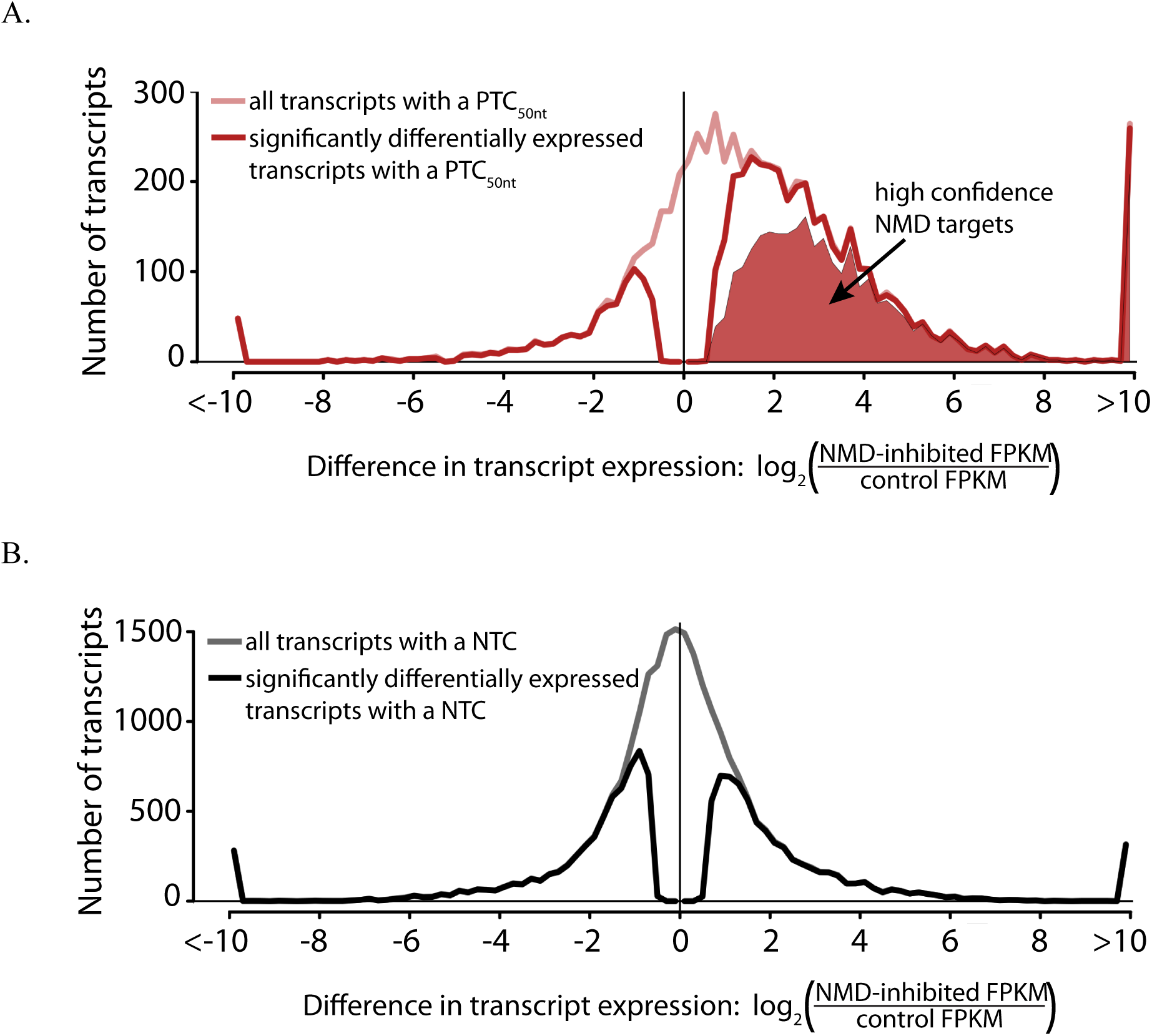
Transcripts with a PTC_50nt_ are more likely to increase in abundance when NMD is inhibited. A) Histogram of expression changes of PTC_50nt_-containing transcripts between NMD inhibited cells and control cells. B) Histogram of expression changes of NTC-containing transcripts between NMD inhibited cells and control cells. Transcripts that were not expressed in one of the conditions were given a log2(inhibited FPKM/control FPKM) value of ±10. PTC_50nt_-containing transcripts demonstrated a strong bias towards accumulation when NMD was inhibited (Kolmogorov-Smirov (KS) test, p < 2.2 × 10-16).

Since UPF1 has multiple functions, including those independent of NMD (Kaygun and Marzluff 2005; Kim et al. 2005; Taylor et al. 2013), its depletion could cause gene expression changes at the transcriptional level. Additionally, mis-regulation of genes regulated via alternative splicing coupled to NMD could have downstream effects. To control for the potential contribution of these transcriptional changes to the increased abundance of PTC_50nt_-containing transcripts, we further focused on PTC_50nt_-containing transcripts that not only were significantly 1.5x more abundant in NMD inhibited cells (and increased in both biological replicates independently), but also adhere to either of the following criteria: 1) no NTC-containing isoform from the same gene was more than 1.2x more abundant in the inhibited NMD sample, or 2) the PTC_50nt_-containing transcript increased at least 2x more than did the sum of the NTC-containing isoforms from the same gene. Note that these criteria require that the genes also produce an NTC-containing isoform that is expressed in this experiment. This will exclude genes that are constitutive NMD targets such as some selenoproteins (Moriarty, Reddy, and Maquat 1998; Seyedali and Berry 2014) and some of the NMD factors themselves (Huang et al. 2011; Yepiskoposyan et al. 2011).

We thus identified 2,793 isoforms (from 2,116 genes) as our high confidence set of direct NMD targets (**Table S2**). These isoforms were derived from 19% of expressed, protein-coding genes, and 22% (2,116 out of 9,440) of alternatively spliced genes in this cell type (alternatively spliced here defined as producing 2 or more expressed isoforms, including the NMD targets). This set of high-confidence NMD targets makes up over 5% of all CDS-containing transcripts produced. While amongst these certainly remain a small number of artifacts, we believe a much larger number of true NMD targets are excluded. Before degradation, many of these NMD-targeted transcripts are produced at levels comparable to those of NTC-containing transcripts (**Figure S2**), and all are expressed at a substantial level, suggesting that they are not merely splicing noise but instead play an important role in regulation. To confirm the expression changes reported by our analysis of the RNA-Seq data, 48 high-confidence NMD-targeted transcripts were checked by qPCR and all had >1.5x increase when NMD was inhibited, consistent with our RNA-Seq results (**Figure S3**).

### Diverse categories of genes are affected by NMD, particularly splicing

A number of human genes have been previously reported to produce NMD-targeted transcripts, including genes encoding splicing factors and NMD factors. Many of the genes with previously validated NMD-targeted isoforms were found in our results. Specifically, we found that the majority of the genes encoding SR proteins expressed in our data generated an NMD-targeted isoform (10 out of 11, example in **Figure 2A**), as previously reported (Lareau, Inada, et al. 2007; Ni et al. 2007). The only SR gene without a high-confidence NMD-targeted isoform in our study is *SRSF9*, which was reported as having a PTC-containing cassette exon based on homology to mouse. A closer investigation showed that there were only a few junction reads supporting the poison cassette exon in our present data, preventing the confident assembly of the isoform. We also found that five NMD factors (*UPF2, SMG1, SMG5, SMG6, SMG7*) reported to be auto-regulated (Huang et al. 2011; Yepiskoposyan et al. 2011) all showed significant gene expression increases of 1.6x to 3.3x when UPF1 was knocked down (**Figure S4**), while *UPF3A* and *UPF3B* showed non-significant gene expression change.

**Figure 2.**
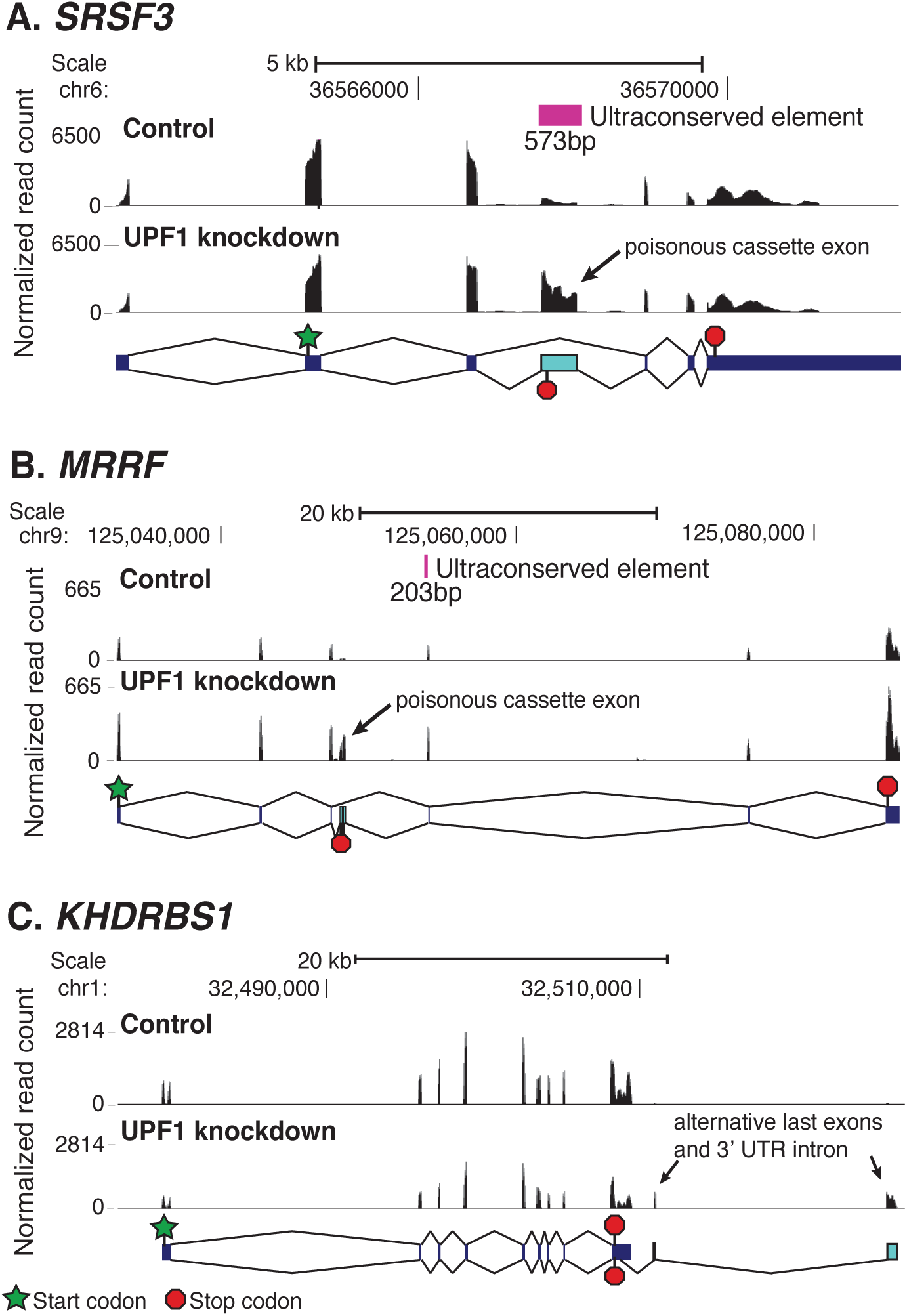
Previously inferred splicing events resulting in NMD targets were confirmed and new ones were discovered (SRSF3, MRRF, KHDRBS1). A) The splicing factor SRSF3 is a previously known NMD target with an ultraconserved element overlapping the PTC_50nt_+ exon. B) MRRF, a mitochondrial ribosome recycling factor, is an example of a non-splicing factor gene with an NMD-targeted transcript associated with an ultraconserved element. The element overlaps the 3’ splice site of the alternative splicing event that generates the PTC_50nt_. C) KHDRBS1, encoding a protein involved in both signal transduction and mRNA processing, has an NMD-targeted transcript generated by alternative splicing and poly-adenylation downstream of the normal termination codon. This figure was generated with the UCSC Genome Brower. Normalized read counts (# reads/10 million mapped reads) for control cells (top track) and NMD-inhibited cells (bottom track) were plotted. Isoform structures are shown below the read tracks. The stop indicated underneath the exon is the PTC_50nt_. A magenta box above the read tracks indicates the presence of an ultraconserved element.

Expanding on the set of splicing-related genes producing NMD-targeted isoforms, we found that a total of 139 genes annotated with the Gene Ontology (GO) term (Ashburner et al. 2000) ‘RNA binding’ and 73 genes annotated with ‘RNA splicing’ produce high-confidence NMD targets (**Table 2**). These GO categories are significantly enriched in NMD-targeted genes (Fisher’s exact test, FDR<0.05, **Figure 3**), as is the KEGG pathway ‘spliceosome’, the SMART domain ‘RNA recognition motif’, and the Uniprot keywords ‘alternative splicing’ and ‘mrna splicng’ (DAVID v6.7 (Huang et al. 2007), FDR<0.10, **Figure S5**). Additionally, the GO categories ‘mitochondrion,’ ‘methyltransferase activity,’ and ‘metabolic process’ are significantly enriched (**Figure 3**) as are the KEGG pathways ‘Aminoacyl-tRNA biosynthesis’, and the Uniprot keywords ‘mitochondrion’, ‘transit peptide’, and ‘polymorphism’ (**Figure S5**). While not significantly enriched, we found that over 4,000 different GO categories contained genes generating NMD-targeted transcripts, indicating NMD-targeted genes are involved in a number of diverse functions such as protein folding, translation, oxidation-reduction processes, vesicle-mediated transport, and chromatin modification. Interestingly, six out of the eight genes annotated ‘NADPH binding’ are high-confidence NMD targets. Categories significantly depleted for genes producing NMD-targeted isoforms include ‘extracellular region’ and some associated with transcription regulation (‘sequence-specific DNA binding’, ‘regulation of transcription, DNA-dependent’, and ‘sequence-specific DNA binding transcription factor activity’).

**Table 2.**
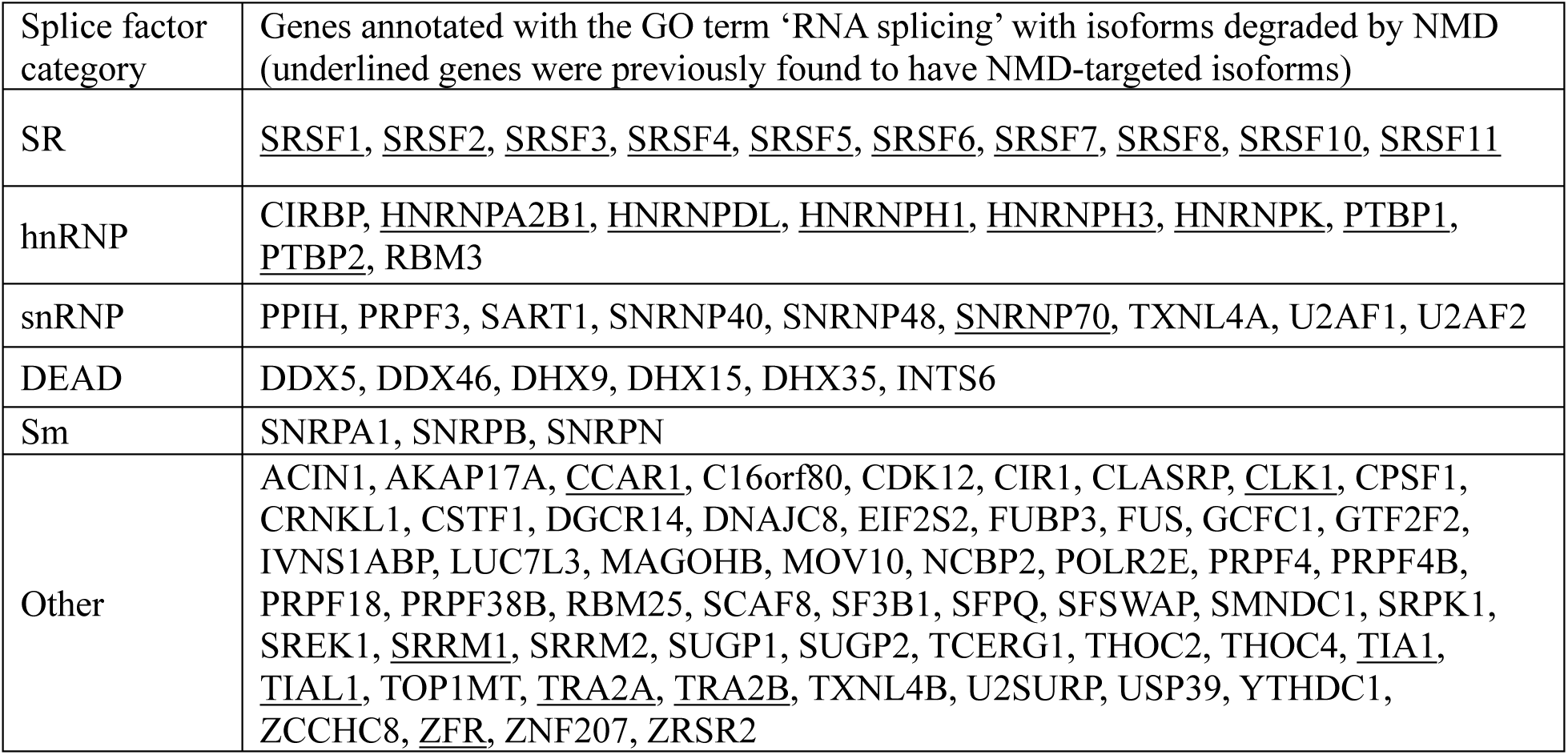
Genes generating an isoform degraded by NMD are enriched for those involved in RNA splicing.

**Figure 3.**
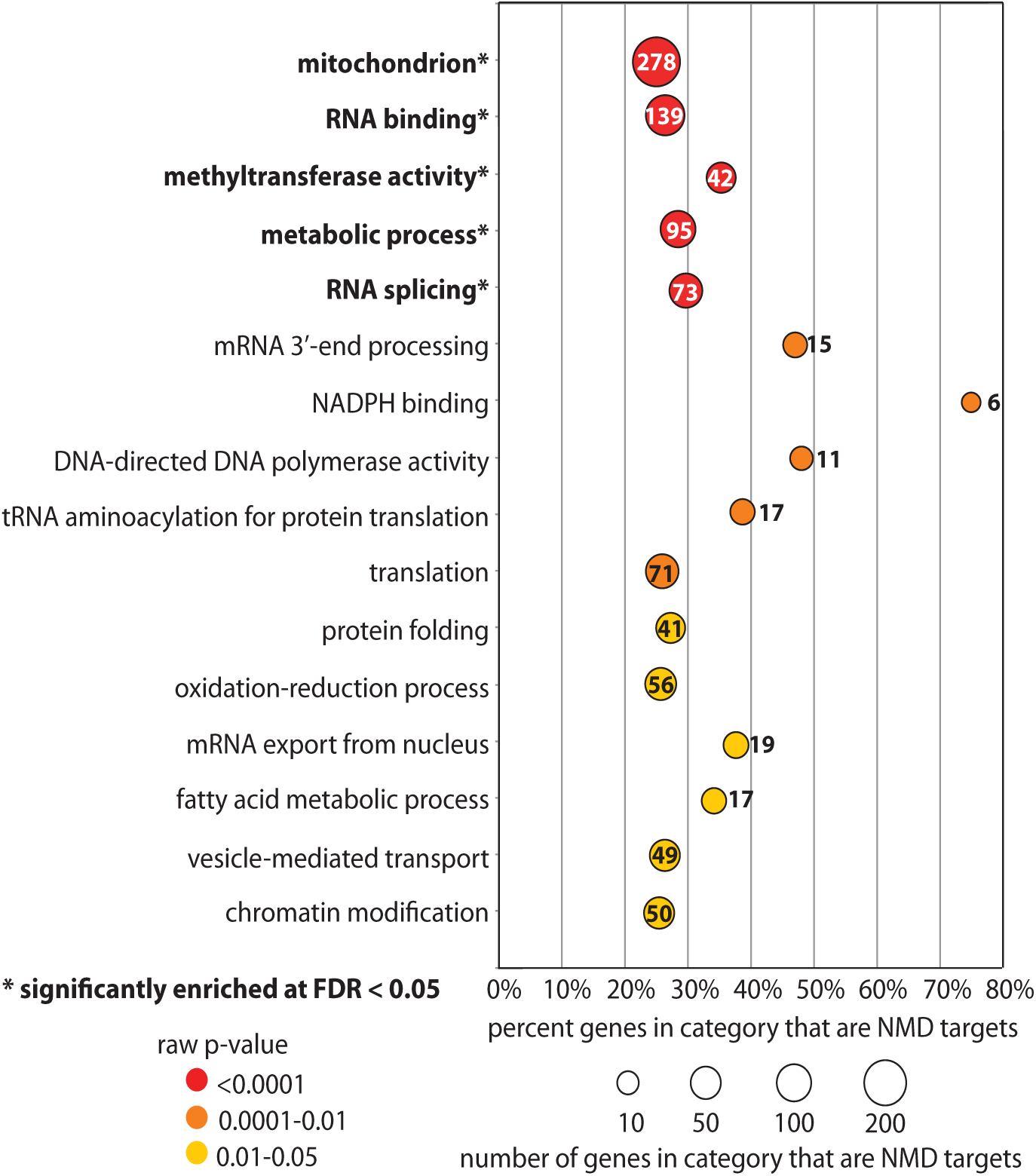
Genes generating an isoform degraded by NMD fall into diverse functional categories. Genes with an isoform degraded by NMD were classified into Gene Ontology functional categories. The asterisk indicates the category is significantly enriched for genes with NMD-targeted isoforms (Fisher’s exact test FDR<0.05; background is genes with an expressed isoform (FPKM>1 in at least one sample)). The test was performed with the fisher.test function in R, the multiple hypothesis correction done in Excel and the plot with ggplot and Adobe Illustrator).

We further found that 71 translation-related genes had NMD-targeted transcripts, including many genes encoding ribosomal proteins (32). This is likely an underestimation of the number of ribosomal genes that are targeted by NMD since most ribosome genes are transcriptionally down-regulated in our NMD-inhibited cells (130 of 169 genes annotated ‘ribosome’ are >1.2x down) and this may mask the increased abundance of NMD-targeted isoforms. In total, over a third of expressed ribosomal genes (58 of 169) produced an alternative isoform with a PTC_50nt_ and are, therefore, potential NMD targets, even if they do not fall into our high-confidence set.

We then investigated the importance of NMD targets in the gene regulatory networks of 35 human tissues published by GTEx (Ardlie et al. 2015). The networks included ∼10,000 protein-coding genes, of which 6,560 were expressed in the Hela cell lines in our study and 1,214 are high-confidence NMD targets. We compared the centrality of the 1,214 NMD targeted genes to the other expressed genes. Centrality measurements (degree, betweenness, and Eigenvector) of NMD targeted genes were not lower than the other genes (Wilcoxon tests, p<0.001 for all tissues); for some tissues, NMD targeted genes even displayed significantly higher level of centralities (**Figure S6**), indicating an important role for these NMD-targeted genes in human gene regulation.

### NMD-targeted genes are enriched for ultraconserved elements

The splicing events that generate PTCs in the SR genes are associated with highly conserved nucleotide sequences (Lareau, Inada, et al. 2007; Ni et al. 2007), including those termed ultraconserved elements (UCEs). UCEs are regions of 100% identity between human, mouse, and rat that are at least 200 bp long (Bejerano et al. 2004). To determine if ultraconserved sequences are more generally enriched in genes producing NMD-targeted isoforms, we examined the overlap between 481 reported UCEs and our high-confidence set of NMD targets. We found that 205 UCEs overlapped 125 expressed genes in our data (83 of which overlapped an exon). Of the 2,116 genes producing an NMD-targeted isoform, 26 overlapped an exonic UCE and 12 genes overlapped an intronic UCE. NMD targets are significantly enriched for UCEs overlapping an exon (Fisher’s exact test, p = 6.7 × 10^−4^) but not for purely intronic UCEs (Fisher’s exact test, p = 0.41).

Intriguingly, for at least 21 of the 26 genes with exonic UCEs, the UCE covers the alternatively spliced region that generates the PTC, supporting the hypothesis that UCEs may be involved in splicing regulation. Most of these alternative splicing events are cassette exons. Interestingly, we found that the majority of the NMD-targeted genes associated with UCEs encode RNA-binding proteins involved in pre-mRNA processing, but those with other functions such as signaling and transcriptional regulation were also found (**Table 3**). One typical example is MRRF, a mitochondrial ribosome recycling factor involved in ribosome release at translation termination, whose alternative isoform has a multiple cassette exon inclusion event that generates a PTC. The downstream exon and relevant intron overlap 201nt of 100% conservation between human and rodent (**Figure 2B**). Some UCEs completely contained within an intron of an NMD-targeted gene may also play a role in regulating splicing to generate PTC-containing isoforms, although the mechanism is less clear.

**Table 3.**
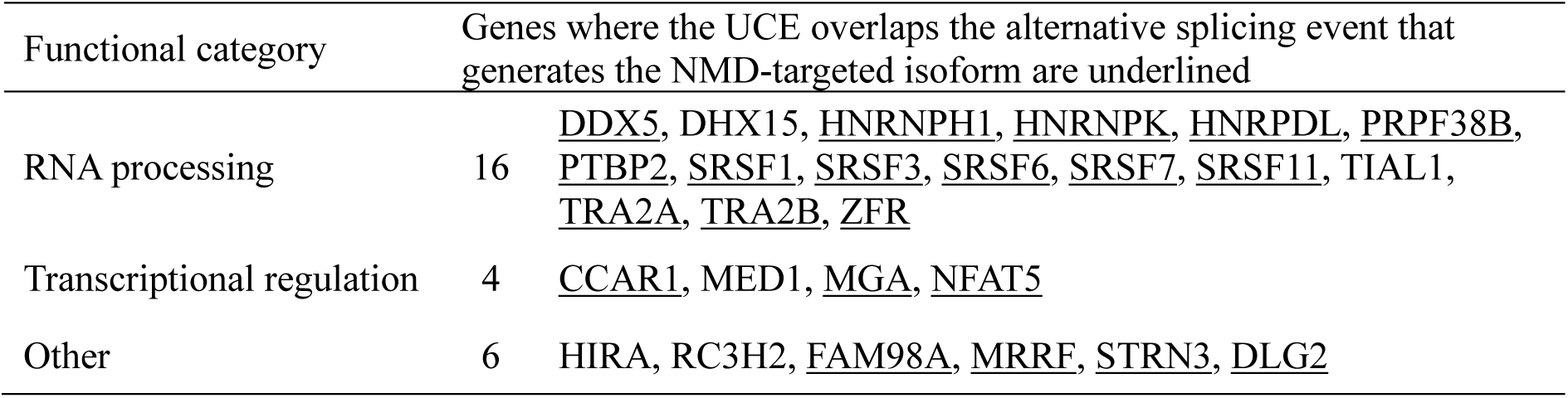
Genes with NMD-targeted isoforms are enriched for ultraconserved elements.

### The 50nt rule is a strong predictor of NMD while a longer 3’ UTR has little effect

While 60% of transcripts containing a PTC_50nt_ were significantly more abundant when NMD was inhibited (and over 70% increased at least 1.2x), a PTC_50nt_ is not believed to be the only trigger of NMD in human cells. The length of the 3’ UTR has also been reported to have an effect on whether a transcript is degraded by NMD. To investigate the correlation between downstream exon-exon junctions (‘50nt rule’) and NMD, we compared the distance between the termination codon and the last exon-exon junction of a transcript with its change in abundance when NMD was inhibited. We found that transcripts with a PTC_50nt_ are twice as likely to increase than those without (**Figure S7A**), and on average they increase more than 3x while on average those without a PTC_50nt_ do not change (**Figures 4A, S9A**). Even for transcripts with a short 3’ UTR (<400 nt), we see a strong correlation of NMD with a downstream exon-exon junction (**Figure 4A inset**). Additionally, there is a clear shift in likelihood of degradation precisely at a distance of 50 nucleotides, as expected by the 50nt rule.

**Figure 4.**
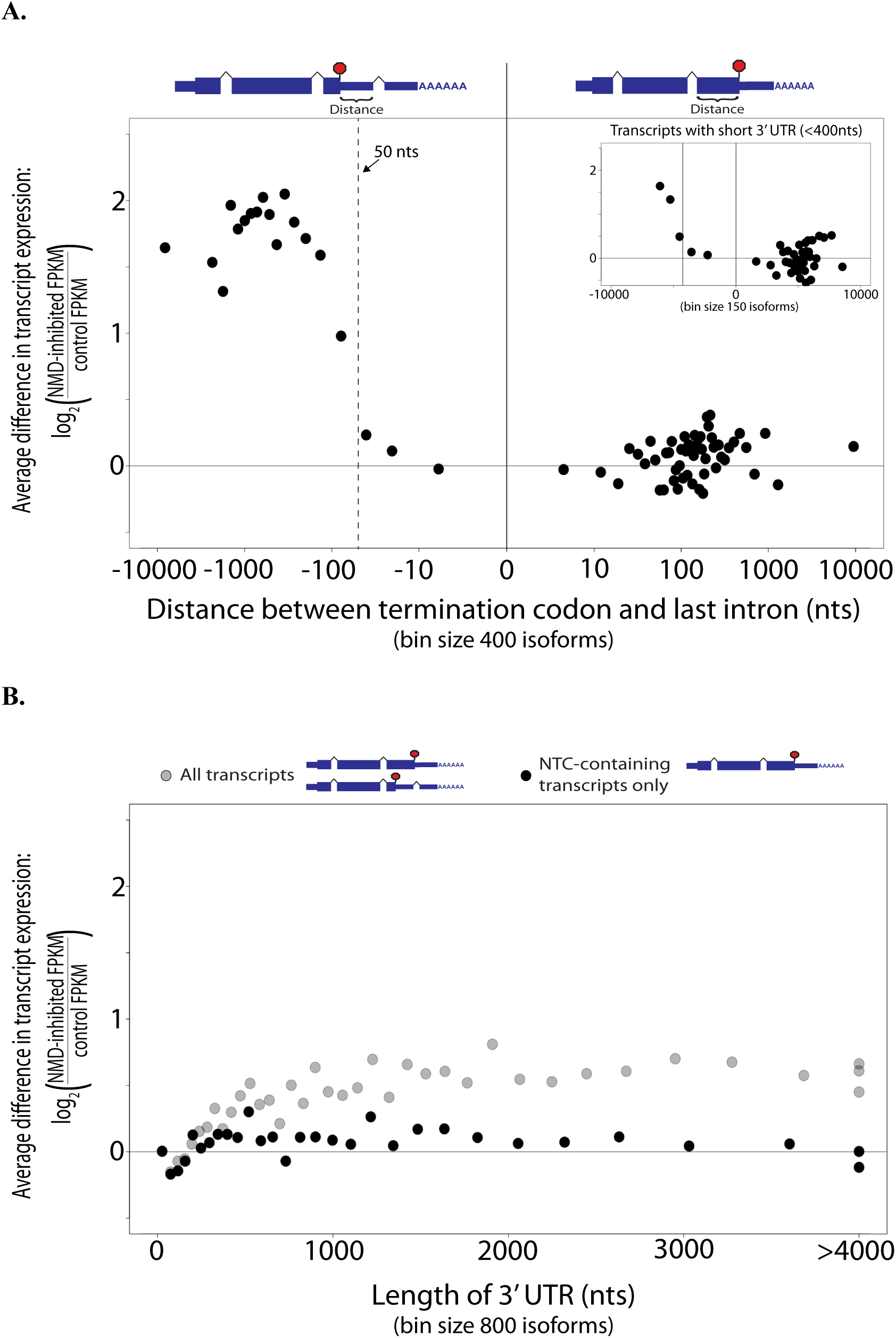
The 50nt rule is a strong predictor of NMD while a longer 3’ UTR is not. A) The distance the termination codon is downstream of the last exon-exon junction versus log2 (inhibited FPKM/control FPKM). Each point is the average log2 (inhibited FPKM/control FPKM) vs the median distance of 400 isoforms. B) Length of the 3’ UTR versus log2 (inhibited FPKM/control FPKM). Gray: all expressed transcripts. Black: Only expressed NTC-containing transcripts. Each point is the average log2 (inhibited FPKM/control FPKM) vs the median distance of 800 isoforms. Bins with median 3’ UTR length more than 4000 nts were placed at 4000 nts. Plots were generated with R.

While we see some correlation between increased 3’ UTR length and increased transcript abundance, this is mostly explained by a strong correlation between 3’ UTR length and the likelihood of a 3’ UTR intron. This correlation largely disappears when looking at only transcripts without the 3’ UTR intron that would create a PTC_50nt_ (**Figures 4B, S7B, S9B**). Without downstream exon-exon junctions, transcripts with a 3’ UTR longer than 2000 nt are significantly more likely to increase in abundance when NMD is inhibited than those with 3’ UTRs shorter than 400 nt (KS test, p = 3.84 × 10^−10^, D = 0.06). However, the effect size is small and the average increase for transcripts with a long 3’ UTR is only 1.02x compared to that for transcripts with a PTC_50nt_, which is 3.3x. A PTC_50nt_ has a strong effect even for transcripts with a 3’UTR less than 400 nt long (2.3x average increase; KS test, p < 2.2 × 10^−16^, D = 0.23). We also investigated the effects of CDS length and GC content of the 3’ UTR. For NTC-containing isoforms, ones with a CDS longer than 2000 nt are significantly more likely to increase in abundance than those with a CDS shorter than 400 nt (KS test, p < 1.58 × 10^−7^, D = 0.07) (**Figures S7C, S8A, S9C**). Similarly, NTC-containing isoforms with 3’ UTR GC content higher than 55% are more likely to increase than those with GC content less than 35% (KS test, p < 2.2 × 10^−16^, D = 0.18) (**Figures S7D, S8B, S9D**).

### Premature termination codons can be generated through alternative splicing events or translation of uORFs

We also explored the types of molecular events that can generate a putative PTC, including alternative splicing and upstream open reading frames (uORFs). Since all of the transcripts in our high-confidence set of NMD targets come from genes that also produce NTC-containing isoforms, we infer that the PTCs were generated by an alternative splicing event. Using the JuncBASE program (Brooks et al. 2011) to investigate these alternative splicing events, we found that at least 365 of the isoforms (13%, of 2,793) were generated by the introduction of a PTC-containing cassette exon (poisonous exon) and 54 isoforms had a retained intron containing a PTC. Additionally, 257 NMD-targeted isoforms (9%) were generated by the introduction of an intron downstream of the termination codon of the productive coding sequence. One interesting example of this is *KHDRBS1*, encoding a protein involved in both signal transduction and mRNA processing, for which alternative splicing combined with alternative poly-adenylation results in the formation of an NMD target without affecting the structure of the coding sequence (**Figure 2C**).

NMD may also act on transcripts with translated uORFs since the uORF termination codon will almost always be recognized as premature, provided that the main coding sequence is not also translated via translation re-initiation. We examined NTC-containing transcripts for uORFs and found that 14,348 NTC-containing transcripts had at least one uORF, but only 5,628 of these had a uORF with a strong Kozak signal sequence for translation initiation (Kozak 1991). Transcripts with a ‘strong’ uORF that either overlaps the main CDS or is at least 35 amino acids long are about twice as likely to significantly increase in abundance when NMD is inhibited, compared to those with no uORF (**Table S3**), similar to what has been reported for plants (Nyikó et al. 2009). Transcripts with a shorter ‘strong’ uORF are also more likely to increase, as are those with ‘weak’ uORFs. Thus, our data suggests that hundreds to thousands of transcripts with uORFs are naturally targets of NMD and could be regulated through the potential translation of uORFs.

## Discussion

In this study, through depletion of UPF1 (a key regulator in the NMD pathway) in HeLa cells, we stabilized transcripts that are otherwise degraded by NMD and determined their structure and expression via RNA-Seq analysis. We used an isoform-centric approach in order to establish a high-confidence set of direct NMD targets and genes that have the potential to be regulated via alternative splicing coupled to NMD. For each transcript, we predicted the coding sequence and did not rely on annotated termination codons, which may be inaccurate and do not exist for novel transcripts. For this analysis, we defined PTCs based on the 50nt rule (Nagy and Maquat 1998) since other features that can trigger NMD have been shown to have low predictive value in our study and others (Colombo et al. 2017; Lindeboom, Supek, and Lehner 2016). While other similar studies use gene-level expression changes to find putative NMD targets (Colombo et al. 2017; Tani et al. 2012), we used isoform-level expression despite the added uncertainty of assigning reads that map to constitutive sequences to the correct isoforms. Crucially, this allowed us to control for transcriptional increases due to secondary effects, which we believe to be very prevalent in the experiment given the large fraction of transcripts that decrease in abundance. This analysis is conservative and likely to miss NMD targets expressed at a lower level or which did not meet our strict requirements.

We report a set of 2,116 genes as high-confidence NMD targets (19% of genes with at least one expressed isoform). The high abundance of these transcripts in the NMD-inhibited sample indicates that they are unlikely to be produced by random errors (**Figure S2**). However, we cannot know what regulatory role they may have and cannot rule out the possibility that they are nonfunctional transcripts with little or no selective impact. Our results re-identified most well-known NMD targets particularly those derived from the SR genes (Lareau, Inada, et al. 2007; Ni et al. 2007), and discovered many more derived from other genes, including many splicing factors. Numerous genes involved in a variety of other functional categories including aminoacyl-tRNA biosynthesis, vesicle-mediated transport, chromatin modification, and metabolic pathways were also found to produce NMD-targeted transcripts, suggesting that a broad range of biological processes may be affected by NMD (**Figure 3**). Transcripts degraded by NMD are significantly enriched for ultraconserved elements of the mammalian genome, supporting a potential role for these elements in regulation through alternative splicing-coupled NMD. Ultimately, we extrapolate that 20% or more of human genes produce NMD-targeted alternative isoforms and are potentially post-transcriptionally regulated by NMD coupled with alternative splicing.

Many NMD-targeted genes, especially splicing factors, are thought to be auto-regulated through a feedback loop wherein protein abundance affects the relative levels of productive and unproductive transcripts. The model is that the protein abundance directly or indirectly affects splicing of its own pre-mRNA so that when a protein is present at an unnecessarily high level, the splicing apparatus shifts to generating NMD-targeted isoforms. We found that when NMD is inhibited, over 500 NMD-targeted genes have decreased expression of NTC-containing isoforms, apparently shifting splicing toward generating the NMD-targeted isoform and likely results in the lower abundance of those proteins. This group of genes includes many involved in RNA splicing. In fact, over half (18/31) of the NMD-targeted genes with the GO annotation ‘RNA processing’ fall into this group, as do all 6 genes annotated as ‘heterogeneous nuclear ribonucleoprotein complex.’ We also found 180 NMD-targeted genes with decreased expression at both the transcriptional level and the NTC-containing isoform level. These are enriched for genes involved in the ribosome and translation, and their down-regulated expression might be a side effect of the UPF1 knockdown, which results in slowed cell growth compared to control cells. Such down-regulation may be mediated through the concerted effort of transcriptional regulation and alternative splicing coupled with NMD. Thus, hundreds of genes were found with evidence of possible regulation through NMD, indicating such a regulatory system may be much more prevalent than previous thought.

The 50nt rule was the first model proposed to be the general signal for NMD degradation in mammals (Nagy and Maquat 1998), and remains the most prevalent model (most cited) with substantial experimental support. However, a longer 3’-UTR has been proposed to also play a role (Hogg and Goff 2010; Hurt et al. 2013; Yepiskoposyan et al. 2011). Results indicating that a long 3’ UTR is enough to cause NMD have come from experiments done with a particular construct (Hogg and Goff 2010), or by looking at the annotated 3’ UTR length of genes or transcripts that increase in overall abundance when NMD is inhibited (Hurt et al. 2013; Yepiskoposyan et al. 2011). In our present data, we identify the precise isoforms that are degraded by NMD in normal cells, thereby allowing for the discovery of novel instances of PTC_50nt_ as well as directly measuring the 3’ UTR length of a specific isoform. Strikingly, the average isoform abundance change when NMD is inhibited is a 3.3x increase for PTC_50nt_-containing transcripts, compared to no change on average for NTC-containing transcripts. Since we find that the majority of PTC_50nt_s fall within the main coding region, those transcripts are going to have longer 3’ UTRs, but a long 3’ UTR in the absence of a PTC_50nt_ has a much smaller effect on isoform abundance. Similar results were recently reported after examining transcript abundance across the different polysome fractions, which can indicate translation and NMD status (Lloyd, et al. 2020). One explanation for this lack of a strong effect by 3’ UTR length on average is that elements in the 3’ UTR near the termination codon have been reported to protect against NMD (Ge et al. 2016; Kishor, Ge, et al. 2019; Toma et al. 2015).

Thirty percent of transcripts with a PTC_50nt_ do not increase at least 1.2x when NMD is inhibited. However, it is possible that as few as 5% are actually escaping NMD. For the others, we see evidence of transcriptional or splicing regulation that could mask the stabilization of the transcript (including transcriptional down-regulation or a shift in splicing toward the productive isoform) and/or aspects that lead to increased uncertainty in the analysis (such as low sequencing coverage, incorrect transcript assembly or CDS definition, and complex alternative splicing patterns). PTC_50nt_-containing transcripts can escape NMD through secondary structure in the 3’ UTR (Eberle et al. 2008), alternative poly-adenylation sites (Gilat and Shweiki 2007), or differential deposition of the exon junction complex required for NMD (Alexandrov et al. 2012; Saulière et al. 2010), as well as other mechanisms (Dyle et al. 2019; Hoek et al. 2019; Kishor, Fritz, et al. 2019; Neu-Yilik et al. 2011; Peixeiro et al. 2012). We conclude that while 3’ UTR length may have an effect for a subset of genes, the 50nt rule is the major mechanism of targeting a transcript for NMD in humans with the caveat that our work is on ‘natural’ NMD targets and it is unclear what happens in ‘unnatural’ mutation cases.

Based on the above findings, we would not expect most transcripts without a PTC_50nt_ to be degraded by NMD. While we find that almost 30% of NTC-containing transcripts significantly increase when NMD is inhibited, almost the same number decrease in abundance, leading us to believe that these changes are likely due to secondary effects of the knockdown experiment. We sought features of these increasing transcripts that may indicate they are targeted for NMD. As described above, we find little evidence that 3’ UTR length significantly affects likelihood of degradation. The length of the coding sequence also does not have a strong effect (**Figure S8A**). We see a correlation between 3’ UTR GC content and likelihood of increased abundance consistent with evidence that UPF1 binds G-rich sequences (Hurt et al. 2013) and that GC motifs in the 3’UTR are important for NMD (Imamachi et al. 2016). Finally, we found that for thousands of transcripts, having a uORF with a strong Kozak signal sequence almost doubles the likelihood that they increase in abundance in UPF1-depleted cells, indicating that NMD may also be involved in extensive regulation of gene expression via a mechanism that is sensitive to the translation of uORFs as has been reported for the *ATF5* gene (Hatano et al. 2013). uORF transcripts are also often bound by a single ribosome, suggesting a low translation rate, and is consistent with many being NMD targets (Lloyd et al. 2020).

In conclusion, alternative splicing coupled to NMD has the potential to massively shape the transcriptome of human cells by directly regulating at least 20% of alternatively spliced genes as well as being a crucial component of splice factor regulation, which itself has numerous downstream regulatory effects. Uncertainty in short-read RNA-Seq analysis may result in some false positives in our NMD target set, however, our conservative approach in defining our high-confidence set means there are likely far more targets that are being missed (false negatives). While this analysis was performed for only one cell line, HeLa, which may have non-physiological events, it is likely the number of NMD targets will increase when investigating other cell types. Since NMD-targeted transcripts are often found at very low levels or not at all in normal cells, this type of regulation is often overlooked when studying a gene or pathway. However, the potential regulatory scope of alternative splicing coupled to NMD demonstrates that researchers should take care to take this regulation into account.

## Materials and Methods

### Knockdown of UPF1 by shRNA

HeLa cells were inoculated on plates in Dulbecco’s MEM medium with 0.1 mM non-essential amino acids and 10% fetal bovine serum, incubated at 37°C and 5% CO_2_. Plates at 80% cell confluency were transfected with a mixture of the plasmids pSUPERpuro-hUpf1 I and II or with pSUPERpuro-Scramble (a gift from Oliver Mühlemann’s lab (Bühler et al. 2006; Paillusson et al. 2005)), whose functions were to knockdown UPF1 and act as a mock control, respectively. Transfections were performed using Lipofectamine™ LTX and PLUS™ Reagents (Invitrogen) according to the manufacturer’s protocol, and the following culture according to the published method (Paillusson et al. 2005). The whole cell lysates were prepared in 1% sodium dodecyl sulfate and incubated at 100 °C for 5 min, then centrifuged at 12,000 rpm for 10 min. The extracted total protein was quantified using the micro BCA™protein assay kit (Pierce Company). Knockdown efficiency was validated by Western blot analysis, using β-actin as the control. Odyssey^®^ infrared imaging system (Li-Cor) was used to quantify and compare the UPF1 protein level between UPF1-knockdown and control samples. Total RNA was extracted using the QIAGEN RNeasy^®^ Mini kit according to the manufacturer’s manual. RNA concentration was determined by NanoDrop 2000^®^, and RNA integrity was determined by 1.2% agar gel and BioAnalyzer. NMD inhibition was validated by amplification of two known NMD-degraded transcripts using oligos from (Lareau, Inada, et al. 2007) by real-time PCR with ABI 7500 fast real time PCR system (Applied Biosystems), and data were analyzed using 7500 software v2.04 (Applied Biosystems).

### Preparation of RNA-Seq libraries

Directional and paired-end RNA-Seq libraries were constructed according to the Illumina protocol, with a few changes: The adapters were prepared according to the reported methods (Vigneault, Sismour, and Church 2008). The PCR process to prepare the library was divided in two steps. In the first step, 3 cycles of PCR were performed (according to the protocol) to prepare the library template, and then the library was run on a 2% agarose gel, fragments of desired size were cut out and isolated by QIAquick^®^ Gel Extraction Kit. A second round of PCR (12 cycles) was performed to enrich the library and then it was purified twice with Agencourt AMPure XP kit as suggested by the Illumina protocol. The libraries were then assayed on a Agilent 2100 BioAnalyzer. These RNA-Seq libraries were prepared from cells with inhibited NMD and control cells for two biological replicates. One biological replicate was sequenced on an Illumina GAIIx machine with 80 bp reads (40 million paired-end reads produced) and the other on HiSeq 2000 with 100 bp reads (180 million paired-end reads produced).

### Transcript assembly and abundance quantification

Paired end reads for each library were aligned to the NCBI human RefSeq transcriptome (Pruitt et al. 2009) with Bowtie (Langmead et al. 2009) to determine the average insert size and standard deviation, required as a parameter by TopHat (Trapnell et al. 2009). The reads of each library were then aligned to the human genome (hg19 assembly, Feb. 2009) using TopHat v1.2.0 with default parameters plus the following: --solexa1.3-quals, --library-type fr-secondstrand, -- coverage-search, --allow-indels, --microexon-search, and --butterfly-search (over 80% of reads aligned). Cufflinks v1.0.1 (Roberts et al. 2011; Trapnell et al. 2010) was used to assemble each set of aligned reads into transcripts with the UCSC known transcript set (downloaded Jan. 2011 from http://genome.ucsc.edu/ (Fujita et al. 2011)) as the reference guide, along with the following parameters: --frag-bias-correct, and --multi-read-correct. Cuffcompare (a sub-tool of Cufflinks) was used to merge the resulting sets of assembled transcripts. Each junction was assigned a Shannon entropy score based on offset and depth of spliced reads across all four libraries (Brooks et al. 2011), see Supplementary Methods for details. Transcripts with a junction that had an entropy score <1 and were not present in the reference annotation were filtered out. Cuffdiff (a sub-tool of Cufflinks) was used to quantify and compare transcript abundance (measured by FPKM, Fragments Per Kilobase per Million reads) between the UPF1 knockdown and control samples. For each sample, the reads from two biological replicates were provided. The following parameters were used: --frag-bias-correct and --multi-read-correct. Only transcripts with FPKM > 1 in either the control or UPF1 knockdown sample were used for further analysis. A transcript was called significantly more abundant in the UPF1 knockdown sample if Cuffdiff called it significantly changing and the change was greater than 1.5x. Significantly decreased transcript abundances were determined in the same way (change was greater than 1.5x).

### Determination of NMD targets

For each transcript, the coding sequence (CDS) was determined as described in the Supplementary Methods, based on (Hansen et al. 2009). A coding sequence was defined to terminate in a premature stop codon (PTC_50nt_) if its termination codon was at least 50 nucleotides upstream of the last exon-exon junction of the transcript (50nt rule in mammals). We inferred transcripts to be NMD targets if they met several criteria. The transcripts must have both a PTC_50nt_ and significantly and >1.5x increased abundance in NMD inhibited (UPF1 knockdown) cells. The transcripts must also increase in each biological replicate when analyzed independently and come from a gene with a NTC-containing isoform with FPKM > 0. A subset of these were deemed more reliable, which we termed our high-confidence NMD-targeted transcripts. High-confidence NMD targeted transcripts adhered to either of the following criteria were kept: 1) No NTC-containing isoform from the gene was more than 1.2x higher in the NMD inhibited sample, or 2) the PTC_50nt_-containing isoform increased at least 2x more than the sum of all NTC-containing isoform FPKMs from the gene in NMD inhibited cells.

Functional classification was based on Ensemble gene annotation, using Gene Ontology (GO) terms (Ashburner et al. 2000), downloaded from Ensembl BioMart Nov 2011. Functional enrichment or depletion was determined by a one-tailed Fisher’s Exact test for NMD targeted genes against the background of all genes with an expressed CDS-containing isoform (FPKM > 1 in at least one sample). A False Discovery Rate threshold of 0.05 was used to correct for multiple tests (4,328 tests done for enrichment (GO categories with at least one NMD target), and 9,673 tests for depletion (all GO categories)). Alternative splicing events were characterized with JuncBASE (version 120623) (Brooks et al. 2011). The JuncBASE pipeline was run on each of the two biological replicates independently with default parameters. The Cuffcompare output GTF, filtered for unsupported junctions, was used as the database for defining exons. A strong Kozak signal for a uORF start codon was defined as [A/G]NNaugG[not U] as recommended by the Consensus CDS Project (Harte et al. 2012).

### Validation by real-time PCR

To validate the potential NMD-targeted transcripts found in the present study, we measured their relative mRNA abundance in cells with UPF1 knocked down and control cells using quantitative real-time PCR. Total RNAs were extracted as described above. cDNAs were synthesized from 5 µg total RNA using the SuperScript II first-strand cDNA synthesis system (Invitrogen) according to the manufacturer’s instructions. The isoform-specific primers used are listed in **Table S4** (those for SRSF genes were from (Lareau, Inada, et al. 2007)). Real-time PCRs were performed on Applied Biosystems 7500 Fast Real-Time PCR System using the respective pair of primers designed with Primer Premier 6.0 and Maxima SYBR Green/ROX qPCR Master Mix (Thermo Scientific). PCR reactions were performed in four replicates, and the expression levels were normalized to that of β-actin control gene (NM_001101).

### Data availability

The raw reads are deposited in the NCBI SRA database under the BioProject Accession ID PRJNA294972 (SRA Accession ID SRP063462). Datasets used in this study are available for download at dash.berkeley.edu available at: https://doi.org/10.6078/D1H019

## Acknowledgements

We thank Adam Roberts and Lior Pachter for help with the optimization of Cufflinks. This work was funded by NIH Grant R01 GM071655 to SEB, GW was supported by a Tang Distinguished Scholarship from QB3 at UC Berkeley, CEF was supported by NIH T32 GM007232 and T32 HG000047 training grants, and by the Department of Defense (DoD) through the National Defense Science & Engineering Graduate Fellowship (NDSEG) Program, and JPBL was supported by the Center for RNA Systems Biology (NIH Grant P50 GM102706 to Jamie Cate).

## Author contributions

CEF: designed and performed analysis and wrote manuscript

GW: designed and performed experiments

JPBL: reviewed and edited manuscript

ZH: performed analysis

ANB: designed analysis

SEB: designed experiment and analysis, reviewed and edited manuscript.

SEB reviewed the penultimate version but not the final submitted version due to an accident injury.

