## Supplemental Figures and Tables for "Transcriptome analysis of alternative splicing-coupled nonsense-mediated mRNA decay in human cells reveals broad regulatory potential"

#### Supplementary Figures

**Figure S1.** (A) Validation of UPF1 knockdown efficiency by Western blot and (B) confirmation of NMD inhibition (p2)

**Figure S2.** Distributions of expression levels of NTC-containing transcripts and transcripts degraded by NMD (p3)

**Figure S3.** qPCR validation of NMD targets (p4)

**Figure S4.** Change in gene expression of NMD factors when NMD is inhibited (p5)

**Figure S5.** KEGG, SMART, and Uniprot enrichments for NMD targets (p6)

**Figure S6.** Centrality of NMD targeted genes compared to non-NMD genes in a gene regulation network (p7)

**Figure S7.** Fraction of transcripts with different features that increase with NMD is inhibited (p8)

**Figure S8.** GC content of 3' UTR may affect NMD degradation and CDS length has little effect (p11)

**Figure S9.** Transcript features correlating with fold change when NMD is inhibited (p13)

#### Supplementary Tables

**Table S1.** Overview of Cufflinks-assembled transcripts from the RNA-seq data (p16)

**Table S2.** Gene list of NMD targets (see attached)

**Table S3.** Features of uORF-containing transcripts that are degraded by NMD (p17)

**Table S4.** qPCR validation primers (see attached)

**Figure S1. Validation of UPF1 knockdown efficiency by Western blot and confirmation of NMD inhibition**

**A.**

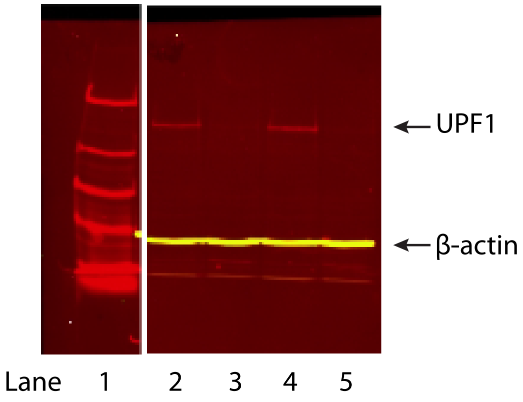


**B.**

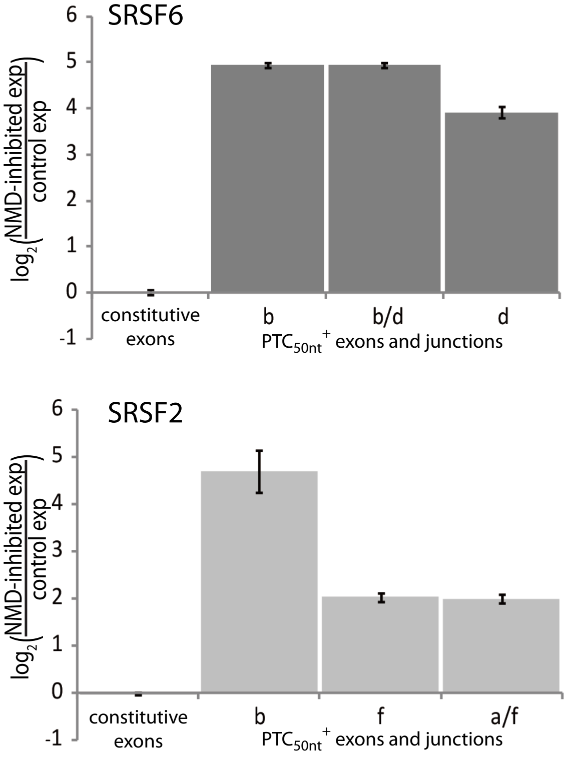


**A.** Western blot evaluating the efficiency of UPF1 knockdown. Lane 1 is the protein ladder; Lanes 2, 4 are the control samples, and Lanes 3, 5 are the UPF1 knockdown samples. **B.** $Log_2$(UPF1-knockdown/control expression) determined by real-time PCR on known NMD-targeted transcripts. Averaged across four biological replicates (two of which were sequenced) and comparing abundance changes for the constitutive exons versus the exons and junctions of the PTC-containing isoforms as previously reported (Lareau et al. 2007).

**Figure S2. Distributions of expression levels of NTC-containing transcripts and high-confidence NMD targeted transcripts**

**A**

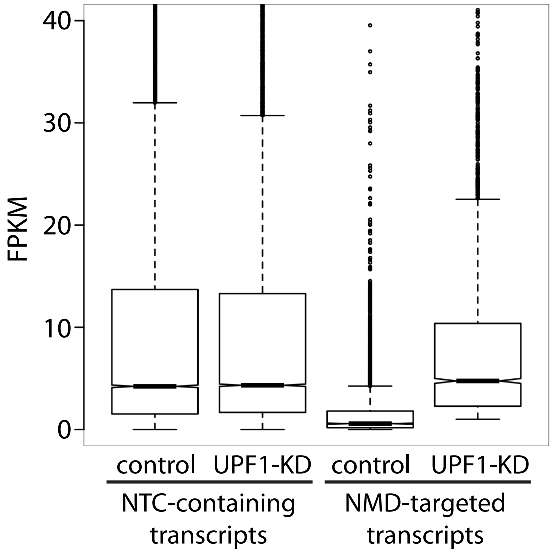


**B**

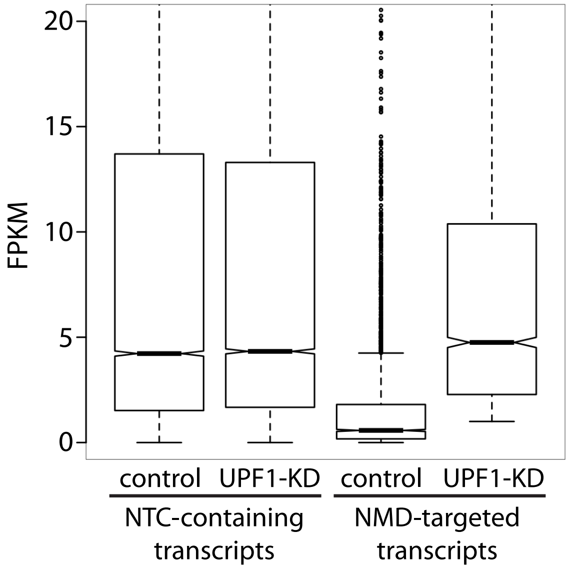


A) Boxplots of the distribution of the expression values (in fragments per kilobase normalized for read depth) for transcripts without a premature termination codon (NTC-containing) and for high-confidence NMD-targeted transcripts in control cells (NMD active) and UPF1-knockdown cells (NMD inhibited). B) Zoomed in version of A. The plots were generated by the R boxplot function and show the median (notch), upper and lower quartiles (box), and ± 1.5 times the interquartile range (whiskers).

**Figure S3.**

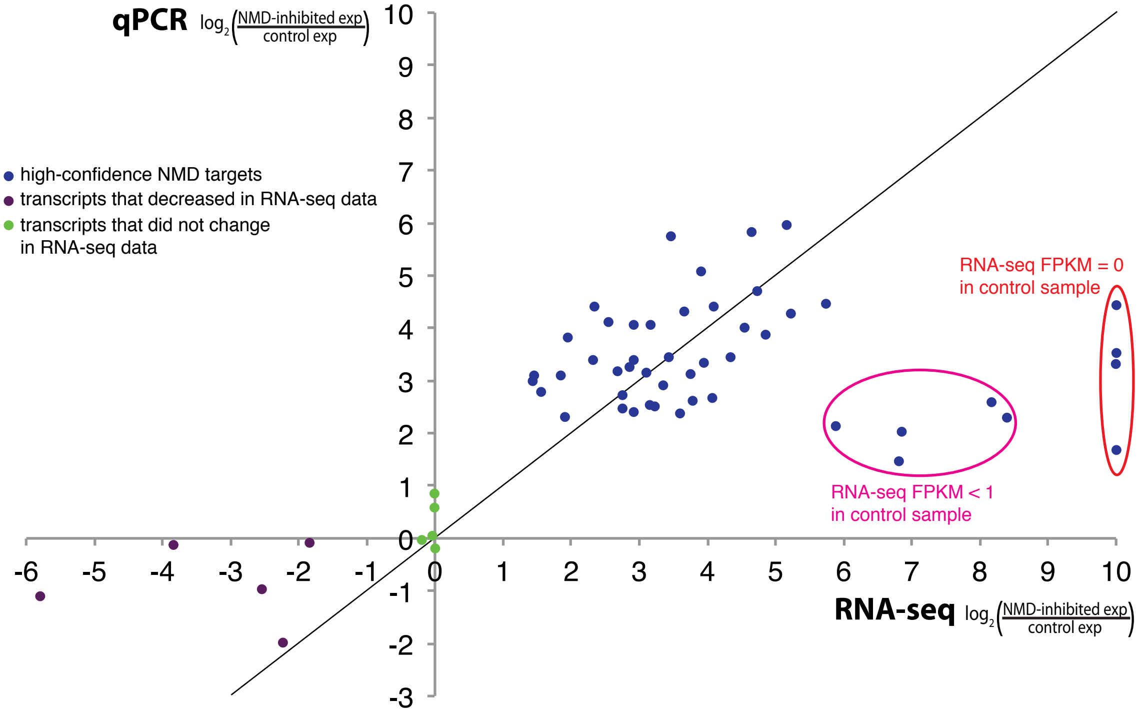


qPCR validation of high-confidence NMD targets. $Log_2$(UPF1-knockdown/control expression) of isoforms in RNA-seq data vs qPCR. Forty-eight high-confidence NMD targets (blue) were tested and all of them increased >2x when measured by qPCR. Note that the four isoforms with a $log_2$(UPF1-knockdown/control expression) of 10 in the RNA-seq data (red circle) were expressed below the level of detection in the control sample (FPKM=0, therefore true fold change is actually infinity but was capped at 10 in this analysis). Additionally, the five isoforms reported to increase substantially more in the RNA-seq data than by qPCR (magenta circle) were expressed at very low levels in the control cells (all had FPKM<1, three had FPKM< 0.2) and this would make it more difficult to accurately quantify their abundance in the control samples and may explain the over-estimation of fold change in RNA-seq data. However, they all still increased >2x when NMD was inhibited using qPCR as measurement. Ten non-NMD targeted isoforms were also measured by qPCR: 5 that did not change in the RNA-seq data (green) and 5 that decreased (purple).

**Figure S4. Change in gene expression of NMD factors when NMD is inhibited**

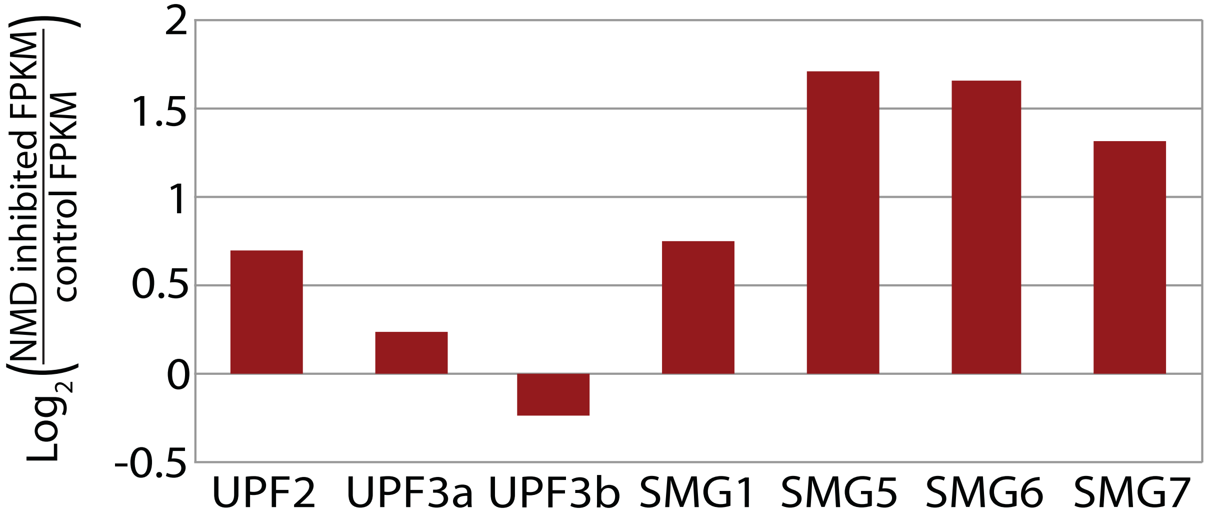


$Log_2$(inhibited FPKM/control FPKM) of gene expression in NMD inhibited cells compared to control cells for NMD factors. *UPF2*, *SMG1*, *SMG5*, *SMG6*, *SMG7* are significantly increased (Cuffdiff, corrected $p<0.05$) (Yepiskoposyan et al. 2011).

#### Figure S5. KEGG, SMART, and Uniprot enrichments for NMD targets

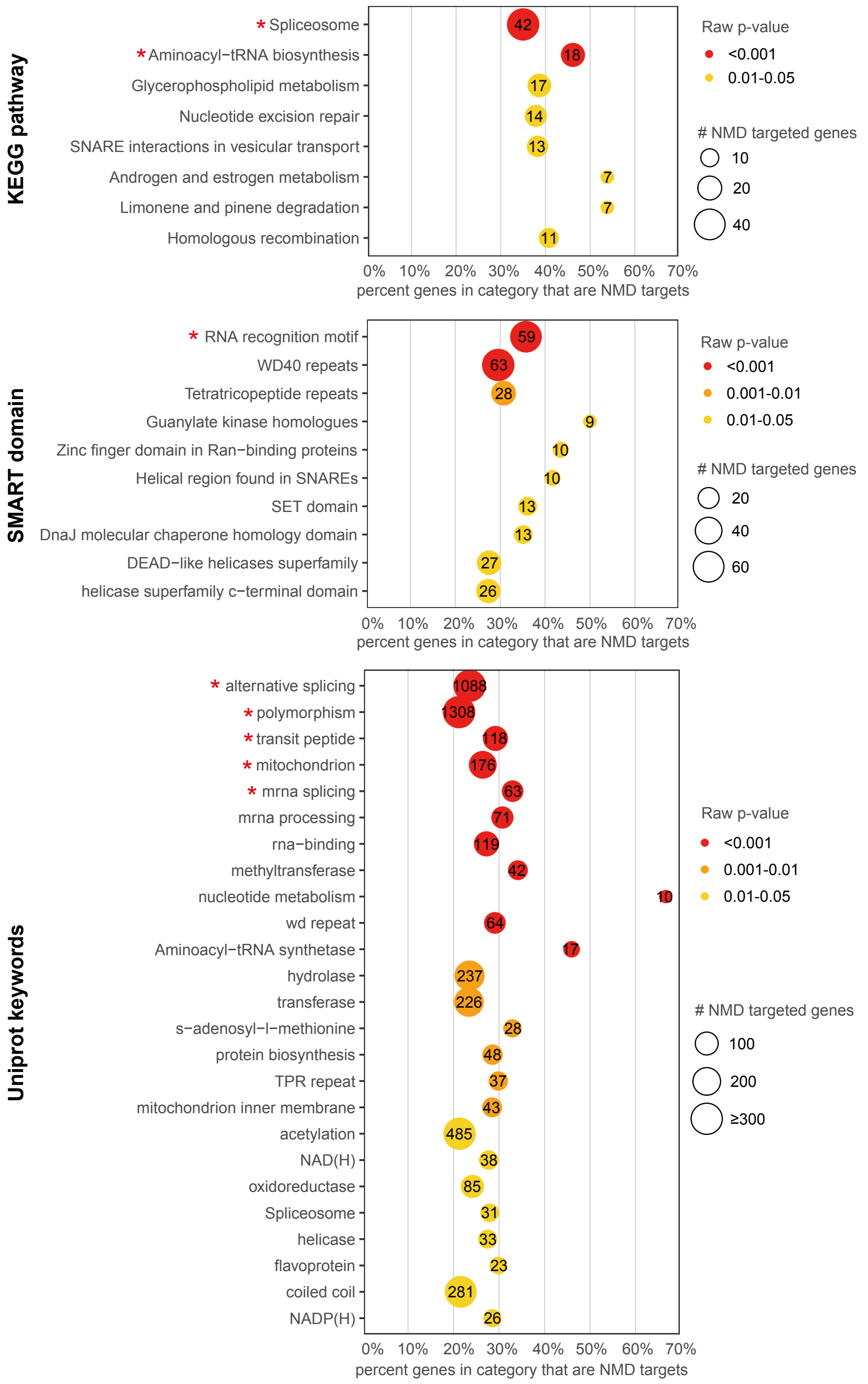


Enrichment analyses were conducted with DAVID v6.7. For all enrichment analyses, 11,056 genes with at least one expressed isoform (FPKM>1 in at least one sample) were used as background. Circle color represents the level of statistical significance and circle size shows number of NMD targeted genes of each category. Red stars represent enriched functions with FDR <0.1.

**Figure S6. Centrality of NMD targeted genes compared to non-NMD genes in a gene regulation network**

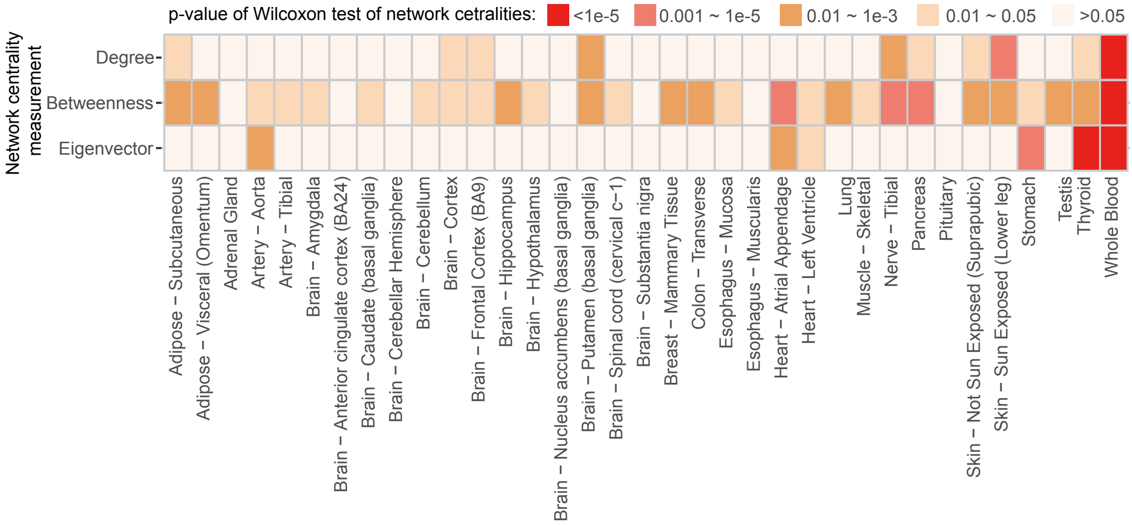


Heatmap illustrating the higher centrality of NMD targeted genes compared to other expressed genes. The p-values of Wilcoxon rank sum tests comparing the centrality of NMD-targeted genes and other expressed genes in GTEX tissue-specific gene expression networks (Ardlie et al. 2015) are plotted. The null hypothesis is that NMD-targeted genes are not more central than other expressed genes and the tests were done on three network centrality measurements: degree, betweenness and Eigenvector.

**Figure S7. Fraction of transcripts with different features that increase with NMD is inhibited**

**A.**

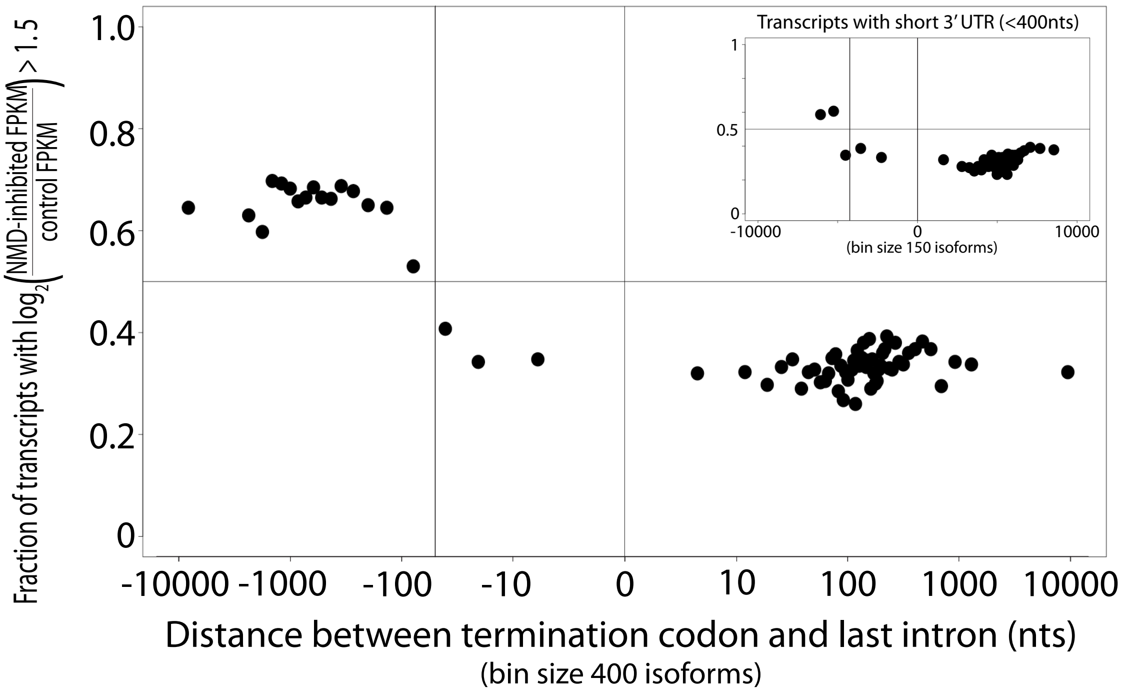


**B.**

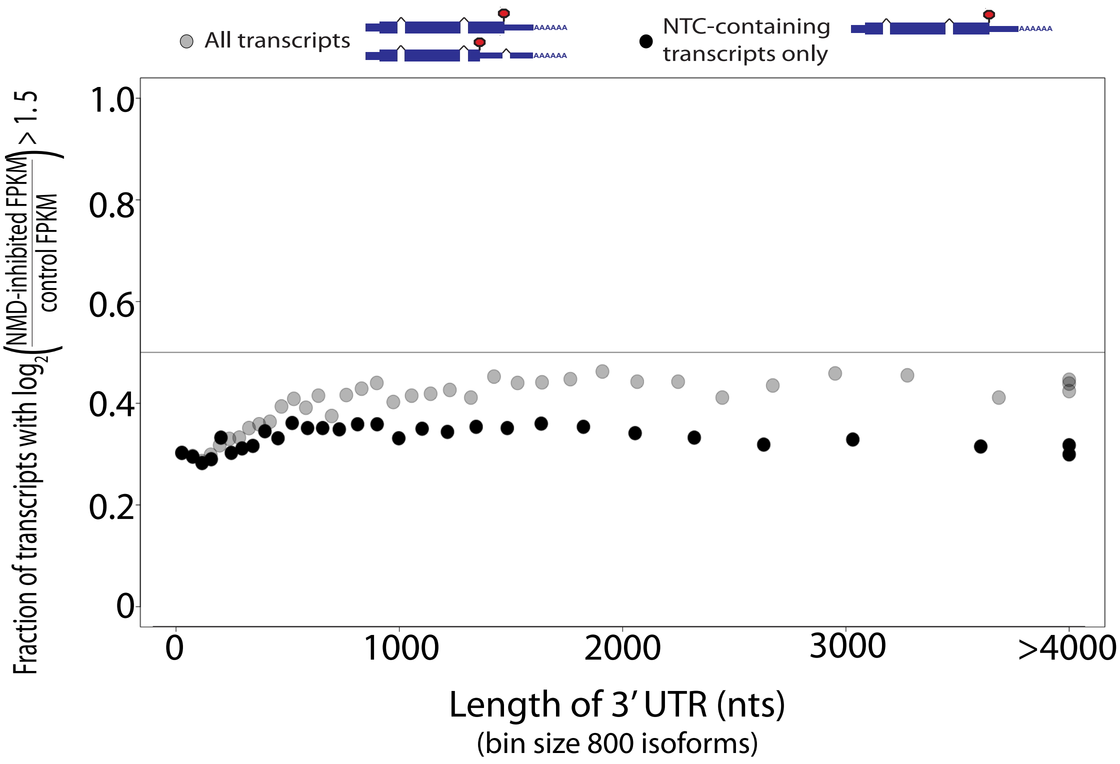

C.

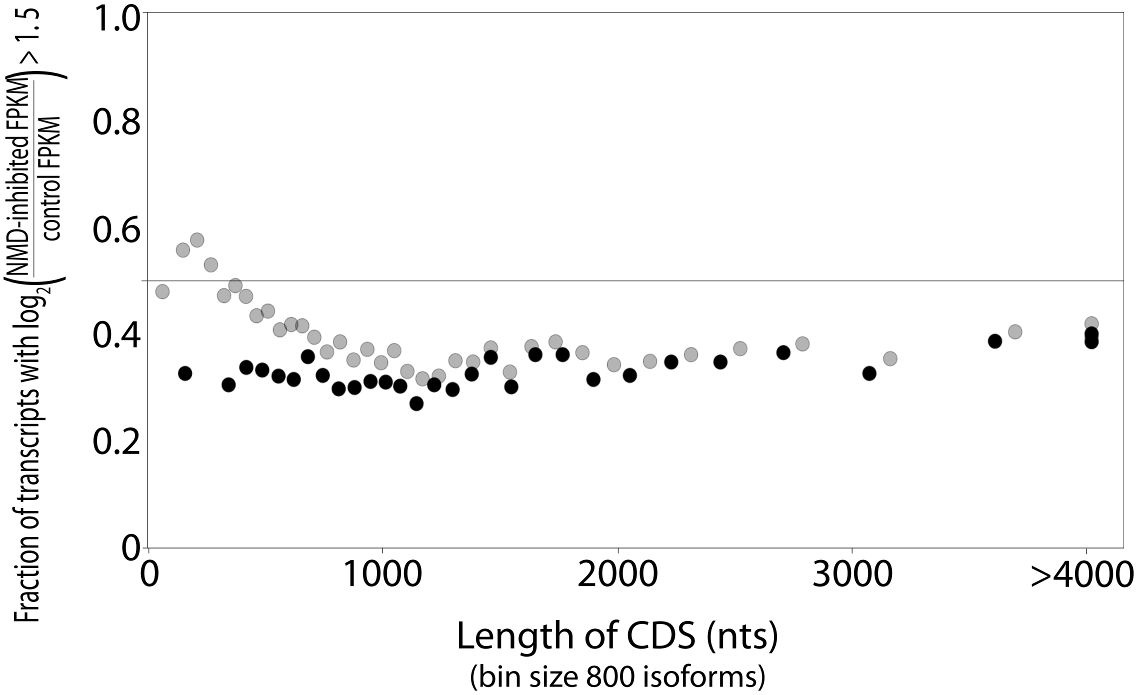

1.0
0.8
0.6
0.4
0.2
0
Fraction of transcripts with $\log_2\left(\frac{\text{NMD-inhibited FPKM}}{\text{control FPKM}}\right) > 1.5$
0
1000
2000
3000
>4000
Length of CDS (nts)
(bin size 800 isoforms)


D.

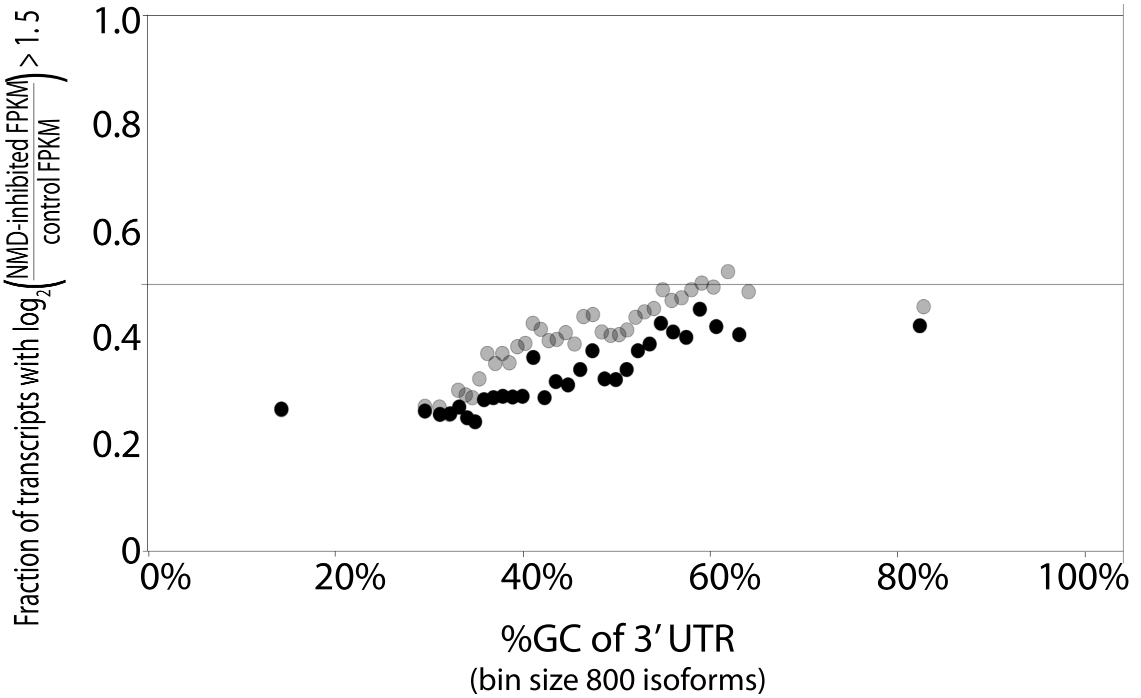

1.0
0.8
0.6
0.4
0.2
0
Fraction of transcripts with $\log_2\left(\frac{\text{NMD-inhibited FPKM}}{\text{control FPKM}}\right) > 1.5$
0%
20%
40%
60%
80%
100%
%GC of 3' UTR
(bin size 800 isoforms)

(A) Distance between stop codon and last exon-exon junction, (B) 3' UTR length, (C) CDS length, or (D) 3' UTR %GC versus the fraction of transcripts that are >1.5x increased in abundance when NMD is inhibited. Gray: all expressed transcripts. Black: Only expressed NTC-containing transcripts. Each point is the average $\log_2$ (inhibited FPKM/control FPKM) vs the median distance or length of 400 or 800 isoforms. Transcripts with a CDS or 3' UTR longer than 4kb fall into the last bin. Plots were generated with R.

**Figure S8. GC content of 3' UTR may affect NMD degradation and CDS length has little effect**

**A.**

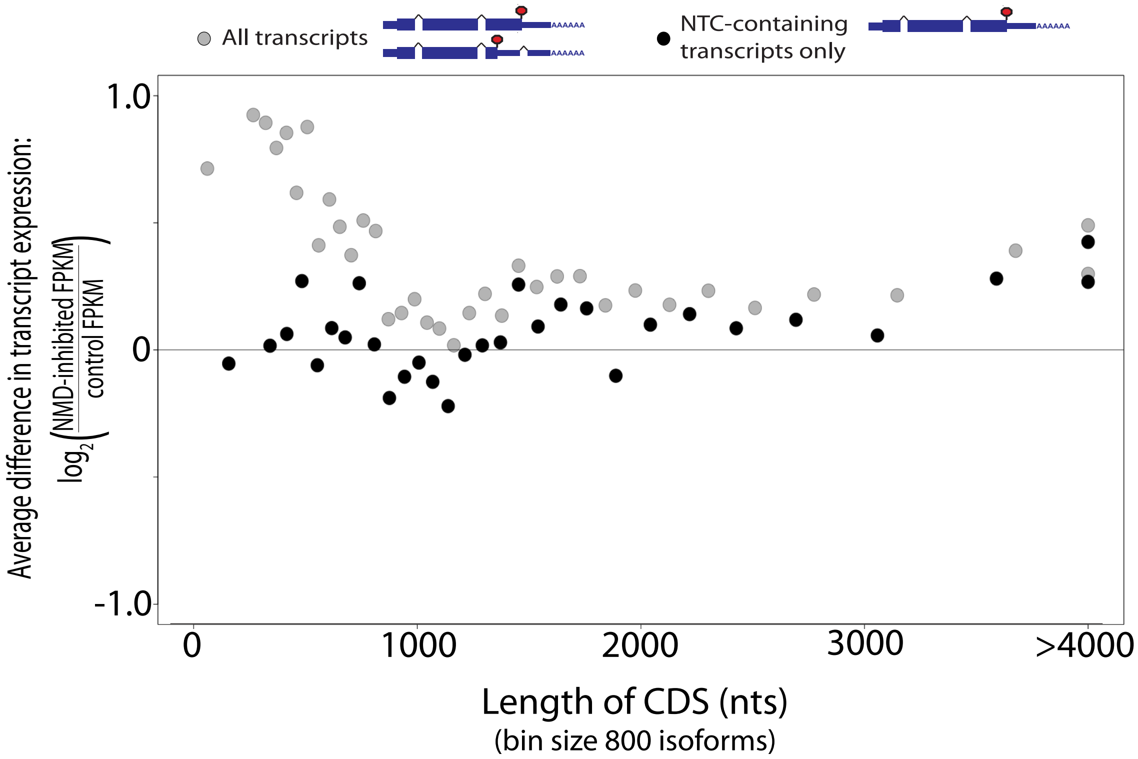


**B.**

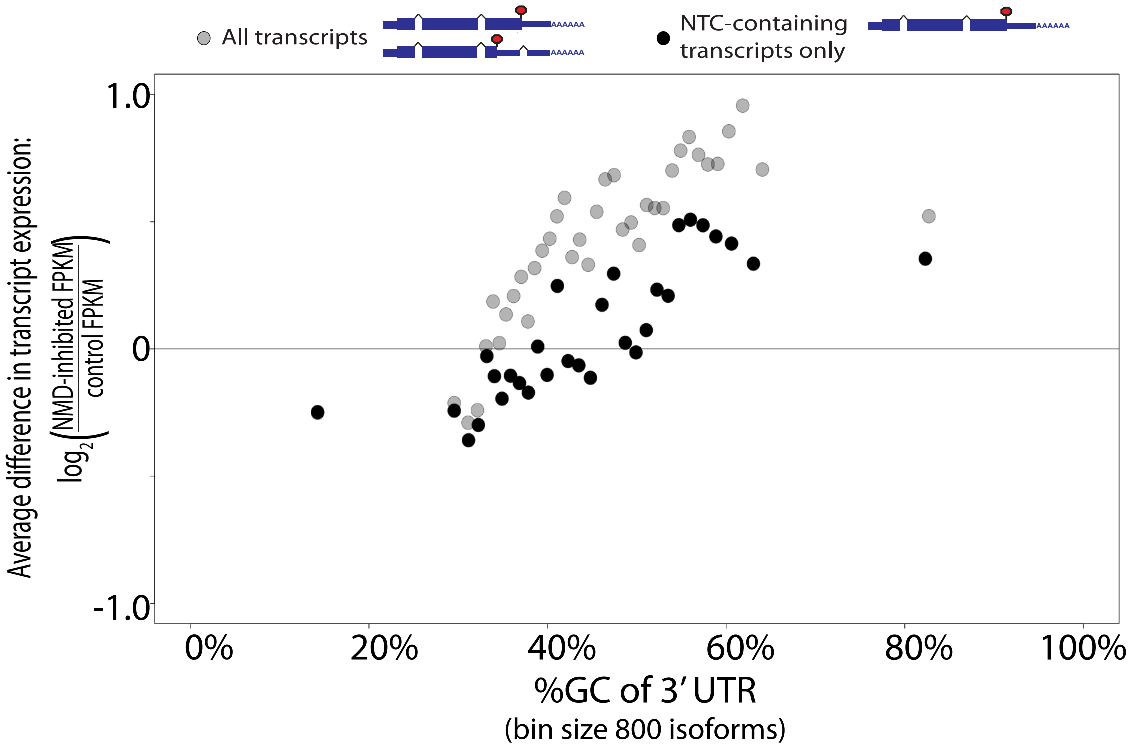

**(A)** CDS length, or (B) 3' UTR %GC versus $log_2$ ratio of expression in NMD inhibited cells to expression in control cells. Gray: all expressed transcripts. Black: Only expressed NTC-containing transcripts. Each point is the average $log_2$ (inhibited FPKM/control FPKM) vs the median of 800 isoforms. Transcripts with a CDS longer than 4kb fall into the last bin. Plots were generated with R.

**Figure S9. Only the distance between the termination codon and the last intron correlates with change in expression when NMD is inhibited**

**A.**

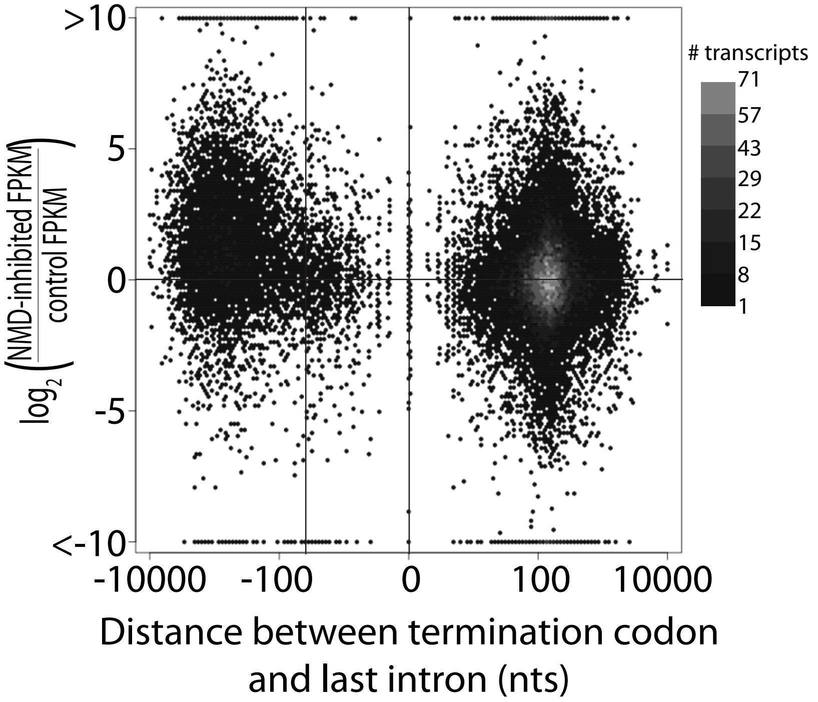


**B.**

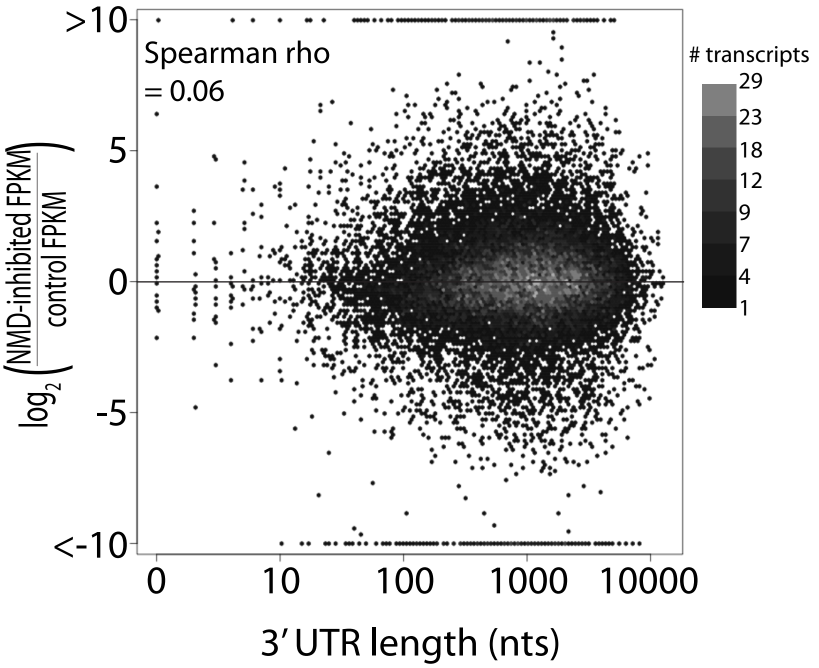

**C.**

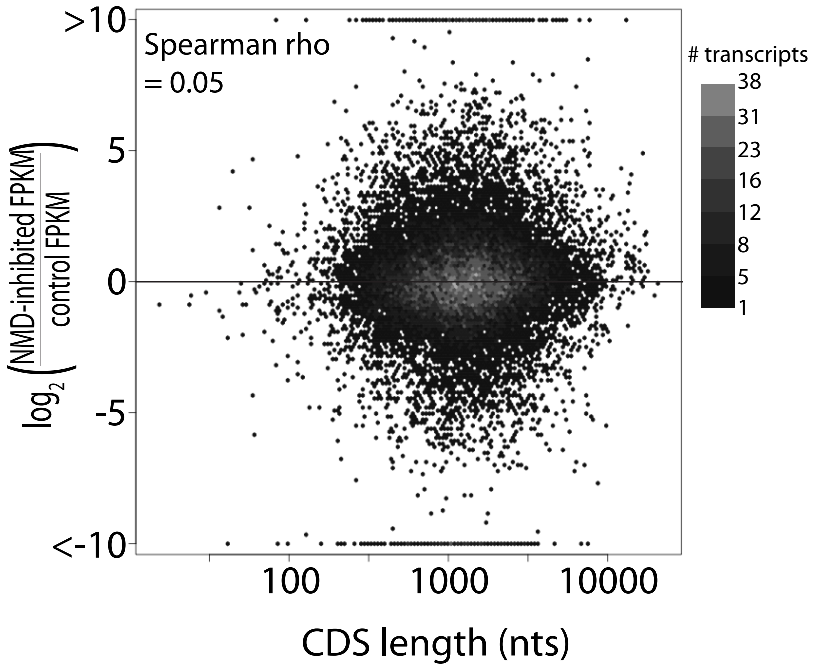
>10
Spearman rho
= 0.05
### transcripts
38
31
23
16
12
8
5
1
5
0
-5
<-10
log2(NMD-inhibited FPKM / control FPKM)
100
1000
10000
CDS length (nts)

**D.**

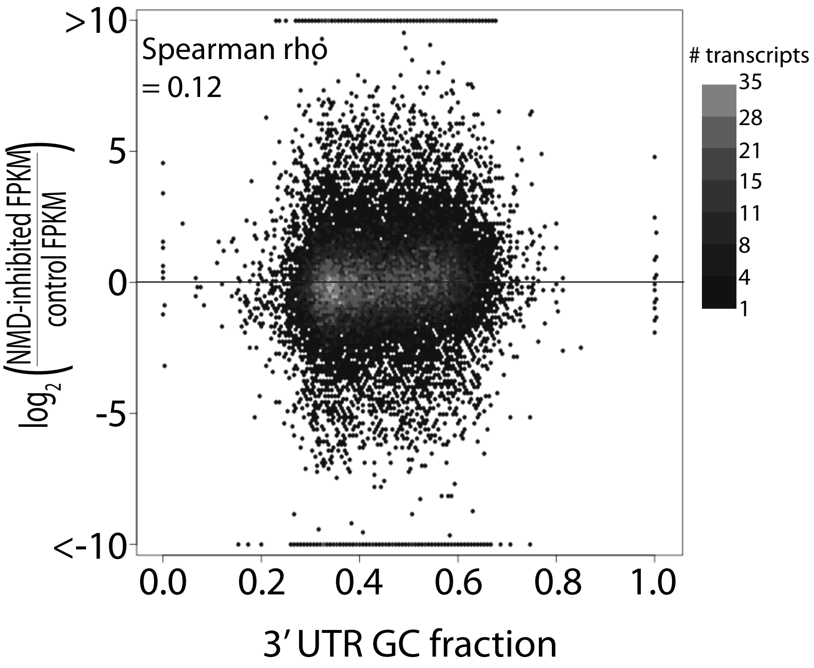
>10
Spearman rho
= 0.12
### transcripts
35
28
21
15
11
8
4
1
5
0
-5
<-10
log2(NMD-inhibited FPKM / control FPKM)
0.0
0.2
0.4
0.6
0.8
1.0
3' UTR GC fraction

(A) Distance between stop codon and last exon-exon junction, (B) 3' UTR length, (C) CDS length, (D) or 3' UTR %GC versus $\log_2$ ratio of expression in NMD inhibited cells to expression in control cells. If the change was > 10x or < -10x, the point was places at 10 or -10, respectively. A, B, C are on a log scale on the x-axis. Lighter coloring indicates multiple transcripts falling into that bin. Plot was made with the hexbin package in R.

**Table S1. Overview of Cufflinks-assembled isoforms from RNA-seq data**

| Class Description | # of isoforms | Expressed isoforms (FPKM >1 in at least 1 sample) | CDS-containing isoforms |
|---|---|---|---|
| Matches reference exactly (=) | 74,233 | 23,687 | 21,616 |
| Contained with reference (c) | 1,627 | 628 | 391 |
| Novel isoform of reference (j) | 16,993 | 11,413 | 8,160 |
| Unknown, intergenic (u) | 16,232 | 10,515 | 0 |
| Probably pre-mRNA (e) | 575 | 279 | 72 |
| Completely intronic (i) | 13,993 | 9,422 | 0 |
| Other generic exon overlap (o) | 467 | 249 | 73 |
| Apparent polymerase run-on (p) | 3,156 | 1,824 | 0 |
| Exon overlap, antisense (x) | 773 | 393 | 0 |
| Intronic, antisense (s) | 0 | 0 | 0 |
| Repeat (r) | 0 | 0 | 0 |
| Multiple classes (.) | 2,954 | 2,565 | 5 |
| TOTAL | 131,003 | 60,976 | 30,317 |

Note: the code in parentheses is assigned by Cufflinks to classify assembled transcript.

**Table S2. Isoforms and gene list of NMD targets.**
See attached text file paper-TableS2.txt

**Table S3. Transcripts with at least one uORF that is long, overlaps the CDS, and/or has a strong Kozak signal are more likely to be degraded by NMD**

| | | | FPKM > 1 | Increased abundance[†] | Decreased abundance† | Ratio[§] |
|---|---|---|---|---|---|---|
| uORF: strong Kozak signal | uORF overlaps main CDS | | 408 | 165 (40%) | 90 (22%) | +1.83 |
| | No overlap | uORF ≥ 35aa | 602 | 269 (45%) | 119 (20%) | +2.26 |
| | | uORF < 35aa | 4,618 | 1,661 (36%) | 1,054 (23%) | +1.58 |
| uORF: weak Kozak signal | uORF overlaps main CDS | | 3,178 | 996 (31%) | 891 (28%) | +1.12 |
| | No overlap | uORF ≥ 35aa | 1,425 | 437 (31%) | 341 (24%) | +1.28 |
| | | uORF < 35aa | 4,117 | 1,113 (27%) | 1,192 (29%) | -1.07 |
| Transcripts with no uORF | | | 9,540 | 2,244 (23%) | 3,041 (32%) | -1.36 |
| All NTC-containing transcripts | | | 23,888 | 6,885 (29%) | 6,728 (28%) | +1.02 |

[†]Increased and decreased abundance refers to the expression changes in NMD-inhibited cells (compared to control cells) that are significant according to Cuffdiff and at least >1.5x different.

[§]A positive ratio (+) is increased/decreased and a negative ratio (-) is decreased/increased. A strong Kozak signal for a uORF start codon was defined as [A/G]NNaugG[not U] as recommended by the Consensus CDS Project (Harte et al. 2012).

**Table S4. Primers for qPCR validation of NMD targets**

See attached excel file paper-TableS4.xls

**References**

Ardlie, Kristin G., David S. DeLuca, Ayellet V. Segrè, Timothy J. Sullivan, Taylor R. Young, Ellen T. Gelfand, Casandra A. Trowbridge, Julian B. Maller, Taru Tukiainen, Monkol Lek, Lucas D. Ward, Pouya Kheradpour, Benjamin Iriarte, Yan Meng, Cameron D. Palmer, Tõnu Esko, Wendy Winckler, Joel N. Hirschhorn, Manolis Kellis, Daniel G. MacArthur, Gad Getz, Andrey A. Shabalin, Gen Li, Yi Hui Zhou, Andrew B. Nobel, Ivan Rusyn, Fred A. Wright, Tuuli Lappalainen, Pedro G. Ferreira, Halit Ongen, Manuel A. Rivas, Alexis Battle, Sara Mostafavi, Jean Monlong, Michael Sammeth, Marta Melé, Ferran Reverter, Jakob M. Goldmann, Daphne Koller, Roderic Guigó, Mark I. McCarthy, Emmanouil T. Dermitzakis, Eric R. Gamazon, Hae Kyung Im, Anuar Konkashbaev, Dan L. Nicolae, Nancy J. Cox, Timothée Flutre, Xiaoquan Wen, Matthew Stephens, Jonathan K. Pritchard, Zhidong Tu, Bin Zhang, Tao Huang, Quan Long, Luan Lin, Jialiang Yang, Jun Zhu, Jun Liu, Amanda Brown, Bernadette Mestichelli, Denee Tidwell, Edmund Lo, Michael Salvatore, Saboor Shad, Jeffrey A. Thomas, John T. Lonsdale, Michael T. Moser, Bryan M. Gillard, Ellen Karasik, Kimberly Ramsey, Christopher Choi, Barbara A. Foster, John Syron, Johnell Fleming, Harold Magazine, Rick Hasz, Gary D. Walters, Jason P. Bridge, Mark Miklos, Susan Sullivan, Laura K. Barker, Heather M. Traino, Maghboeba Mosavel, Laura A. Siminoff, Dana R. Valley, Daniel C. Rohrer, Scott D. Jewell, Philip A. Branton, Leslie H. Sobin, Mary Barcus, Liqun Qi, Jeffrey McLean, Pushpa Hariharan, Ki Sung Um, Shenpei Wu, David Tabor, Charles Shive, Anna M. Smith, Stephen A. Buia, Anita H. Undale, Karna L. Robinson, Nancy Roche, Kimberly M. Valentino, Angela Britton, Robin Burges, Debra Bradbury, Kenneth W. Hambright, John Seleski, Greg E. Korzeniewski, Kenyon Erickson, Yvonne Marcus, Jorge Tejada, Mehran Taherian, Chunrong Lu, Margaret Basile, Deborah C. Mash, Simona Volpi, Jeffery P. Struewing, Gary F. Temple, Joy Boyer, Deborah Colantuoni, Roger Little, Susan Koester, Latarsha J. Carithers, Helen M. Moore, Ping Guan, Carolyn Compton, Sherilyn J. Sawyer, Joanne P. Demchok, Jimmie B. Vaught, Chana A. Rabiner, and Lockhart. 2015. "The Genotype-Tissue Expression (GTEx) Pilot Analysis: Multitissue Gene Regulation in Humans." *Science*.

Harte, Rachel A., Catherine M. Farrell, Jane E. Loveland, Marie-Marthe Suner, Laurens Wilming, Bronwen Aken, Daniel Barrell, Adam Frankish, Craig Wallin, Steve Searle, Mark Diekhans, Jennifer Harrow, and Kim D. Pruitt. 2012. "Tracking and Coordinating an International Curation Effort for the CCDS Project." *Database : The Journal of Biological Databases and Curation* 2012:bas008.

Lareau, Liana F., Angela N. Brooks, David A. W. Soergel, Qi Meng, and Steven E. Brenner. 2007. "The Coupling of Alternative Splicing and Nonsense-Mediated MRNA Decay." *Advances in Experimental Medicine and Biology* 623:190–211.

Yepiskoposyan, Hasmik, Florian Aeschimann, Daniel Nilsson, Michal Okoniewski, and Oliver Mühlemann. 2011. "Autoregulation of the Nonsense-Mediated MRNA Decay Pathway in Human Cells." *RNA (New York, N.Y.)* 17(12):2108–18.
