## Supplemental Methods for "Transcriptome analysis of alternative splicing-coupled nonsense-mediated mRNA decay in human cells reveals broad regulatory potential"

### Supplementary Methods

Calculating entropy scores for splice junctions

In order to filter out low-confidence or unsupported novel splice junctions, a Shannon entropy score was calculated for each junction using JuncBASE (Brooks et al. 2011). The equation is as follows:

$$\text{Shannon entropy} = -\sum_{i=1}^{n} p(x_i)\log_2 p(x_i)$$

where $n$ is the number of offsets (read starting positions) for the junction and $p(x_i)$ is the number of reads at offset $i$ divided by the total number of reads crossing the junction. Entropy scores were calculated using all mapped reads from all four libraries (two control samples and two experimental samples).

Determining the coding sequence of each isoform

In order to see if an isoform has a $PTC_{50nt}$, we first had to determine the coding sequence (CDS) for each transcript. We did not merely use the CDS defined by a reference annotation for two reasons: 1) NMD targets are likely to be novel isoforms and therefore not have an annotated CDS and 2) many annotated CDS are computationally predicted and not necessarily accurate. Thus, we determined the CDS for each isoform based on the method described in (Hansen, et al 2009).

We started with the set of start codons found in the 'UCSC Genes' reference annotation from the UCSC Genome Browser (downloaded Jan. 2011 from http://genome.ucsc.edu/). For each isoform, we found each ORF produced by starting at every overlapping annotated start codon. For each gene, we found the longest ORF, with priority given to ORFs from isoforms that did not increase upon UPF1 knockdown (FPKM>0 in at least one of the two conditions and <1.2x increase in abundance). For genes with no isoform that did not increase, the longest ORF from any isoform was chosen. For isoforms with an ORF that had the same start and stop codons as the gene's longest ORF, that ORF was chosen to be the CDS. For isoforms that did not overlap the start codon of the gene's longest ORF, the longest ORF for that isoform was chosen to be the CDS. And for isoforms that overlap the start codon of the gene's longest ORF but have a different stop codon, we chose the longest ORF produced from either that start codon or from an ATG (including those not in the annotation) found upstream of that start codon but not found in the isoforms with the longest ORF start and stop codon.
